# Accurate identification and quantification of commensal microbiota bound by host immunoglobulins

**DOI:** 10.1101/2020.08.19.257501

**Authors:** Matthew A. Jackson, Claire Pearson, Nicholas E. Ilott, Kelsey E. Huus, Ahmed N. Hegazy, Jonathan Webber, B. Brett Finlay, Andrew J. Macpherson, Fiona Powrie, Lilian H. Lam

## Abstract

**Background:** Identifying which taxa are targeted by immunoglobulins can uncover important host-microbe interactions. Immunoglobulin binding of commensal taxa can be assayed by sorting bound bacteria from samples and using amplicon sequencing to determine their taxonomy, a technique most widely applied to study Immunoglobulin A (IgA-Seq). Previous experiments have scored taxon binding in IgA-Seq datasets by comparing abundances in the IgA bound and unbound sorted fractions. However, as these are relative abundances, such scores are influenced by the levels of the other taxa present and represent an abstract combination of these effects. Diversity in the practical approaches of prior studies also warrants benchmarking of the individual stages involved. Here, we provide a detailed description of the design strategy for an optimised IgA-Seq protocol. Combined with a novel scoring method for IgA-Seq datasets that accounts for the aforementioned effects, this platform enables accurate identification and quantification of commensal gut microbiota targeted by host immunoglobulins.

**Results:** Using germ-free and *Rag1*^*−/−*^ mice as negative controls, and a strain-specific IgA antibody as a positive control, we determine optimal reagents and fluorescence activated cell sorting (FACS) parameters for IgA-Seq. Using simulated IgA-Seq data, we show that existing IgA-Seq scoring methods are influenced by pre-sort relative abundances. This has consequences for the interpretation of case-control studies where there are inherent differences in microbiota composition between groups. We show that these effects can be addressed using a novel scoring approach based on posterior probabilities. Finally, we demonstrate the utility of both the IgA-Seq protocol and probability-based scores by examining both novel and published data from *in vivo* disease models.

**Conclusions:** We provide a detailed IgA-Seq protocol to accurately isolate IgA-bound taxa from intestinal samples. Using simulated and experimental data, we demonstrate novel probability-based scores that adjust for the compositional nature of relative abundance data to accurately quantify taxon-level IgA binding. All scoring approaches are made available in the IgAScores R package. These methods should improve the generation and interpretation of IgA-Seq datasets and could be applied to study other immunoglobulins and sample types.

## Background

The immunoglobulin isotype A (IgA) is secreted by plasma cells at mucosal barrier sites such as the gastrointestinal (GI) tract, which is also host to the densest community of commensal bacteria in the human body[1, 2]. Intestinal homeostasis requires tolerance to the microbiota but a robust defence against pathogens, and growing evidence suggests that IgA binding is both taxon-dependent and context-specific. IgA in the mucus layer forms a protective barrier between the epithelium and the luminal microbiota [3–5]. Some commensal bacteria have adapted to this specialized niche to promote homeostasis. *Bacteroides fragilis* produces a surface polysaccharide that promotes IgA binding and facilitates mucus layer colonization [6] and the mucin-degrader *Akkermansia muciniphila* induces host IgA and IgG responses[7]. Other studies have suggested that low-affinity IgA contributes to microbiota maintenance, while high-affinity IgA acts on pathogens to promote clearance and inhibit virulence [8–15]. Pathobionts are members of the microbiota that are typically harmless under homeostatic conditions but can drive disease given certain environmental stimuli[16]. IgA binding of taxa might also reflect such pathogenic potential and IgA bound gut bacteria have been implicated in inflammatory disease [17–20]. Accurate identification of commensal microbiota bound by IgA can therefore offer important insights into the mechanisms underlying host microbial mutualism and its dysregulation in disease.

Microbiota-wide profiling of IgA bound taxa can be achieved using IgA-Seq. Anti-IgA antibodies are used to stain a complex gut microbiota, then bound and unbound bacteria are separated by Fluorescence Activated Cell Sorting (FACS) or Magnetic Activated Cell Sorting (MACS) and identified by 16S rRNA gene sequencing [17, 18]. We refer to the sorted IgA bound bacteria as the IgA+ fraction, the sorted unbound bacteria as the IgA- fraction, and the sample before sorting as the pre-sort sample. IgA-Seq has been used to uncover antibody targeting of taxa by both T-cell dependent and T-cell independent pathways [7, 10, 12, 21, 22], as well as differential binding of taxa during disease. IgA-Seq was first described by Palm et al. to identify taxa differentially bound by IgA in inflammatory bowel disease (IBD) patients [17] and was adapted by Kau et al. to identify increased binding of *Enterobacteriaceae* by IgA in malnourished infants [18]. Variations of these protocols have since been applied to study IgA binding in other disease states, employing a range of IgA-Seq protocols utilising different reagents, sort modalities and configurations [19, 20]. Some of these parameters have been validated individually in prior studies but there has yet to be a comprehensive description of how to design and benchmark an optimal IgA-Seq protocol.

More importantly, there has yet to be a formal consideration of the analytical approaches used to score taxon binding in IgA-Seq datasets. IgA-Seq does not quantify the affinity of IgA antibodies directly, but rather provides a measure of overall IgA binding. The likelihood of bacteria within a given taxon being bound by IgA will be influenced by a combination of the affinities of the IgA pool for the taxon, the spatial distribution of those bacteria, and the expression of relevant epitopes by the taxon. Additionally, the IgA+ abundance does not solely represent this likelihood of taxon binding. It is also a function of the number of bacteria present: a taxon highly targeted by host IgA could make up a low proportion of the IgA+ fraction if only a few bacteria are present and vice versa. The principal aim of IgA-Seq is not to solely identify which taxa are most bound by IgA but rather to identify those taxa with a higher likelihood of binding relative to other bacteria. As a result, indices that account for the initial quantities of bacterial taxa must be applied to quantify IgA-Seq data.

In IgA-Seq studies to date, IgA binding has largely been scored using indices as described by either Palm et al. or Kau et al. (referred to here as the Palm and Kau indices for clarity)[17, 18]. These scores compare the abundance of a taxon in the IgA+ fraction to its abundance in the IgA- fraction. This accounts for the overall pre-sort quantities of taxa on the assumption that, as both fractions are drawn from the same starting pool of bacteria, the IgA+ and IgA- abundances are representative of the proportion of a taxon’s bacteria bound by IgA. However, these scores are influenced by an additional confounder – that these are relative abundances. As a result, an increase in one taxon will inherently reduce the percentage contribution of others. Thus, the IgA+ and IgA- abundances of a given taxon are also influenced by the starting abundances and IgA binding of all other taxa. Scores like the Kau and Palm indices correlate with taxon-level IgA binding but the necessity to use relative abundances in IgA-Seq experiments abstracts their interpretation. We will show that, without careful consideration of these effects, it is possible to misinterpret results observed using these metrics.

Here, we aim to address these methodological and analytical limitations to improve the accessibility, quality and quantitative interpretation of IgA-Seq experiments. Using germ-free and immune-deficient mice as negative controls and a strain-specific IgA antibody for positive controls, we describe the methods that can be used to validate the quality of each stage of an IgA-Seq experiment and derive an optimised IgA-Seq protocol. In lieu of a gold standard for benchmarking IgA binding scores, we develop a platform for simulating an IgA-Seq experiment where the underlying relative IgA binding levels between taxa are known. Using this, we show that existing indices used to score taxon-level IgA binding are influenced by relative abundances, which needs to be taken into account when drawing conclusions. We present novel probability-based scores that circumvent this by providing direct quantification of the proportion of bacteria belonging to a given taxon that are bound by IgA. Finally, we demonstrate the utility of the protocol and novel probability scores by applying them to study IgA binding in mouse models of colitis and malnourishment. These methods facilitate generation of high-quality IgA-Seq data and direct quantification of relative immunoglobulin binding between taxa.

## Results

### Quality control standards for bacterial flow cytometry and IgA staining

We first determined the optimal gating strategy for isolating bacteria from faecal and colonic samples by FACS. The BD LSRFortessa X20 and FACSAria III flow cytometers were selected based on an identical configuration with 405, 488, 633 and an additional 561nm laser, which eliminated PE-FITC spill over and increased PE sensitivity. Whereas eukaryotic cells have diameters ranging from 10–100 microns, bacteria generally range from 0.3–5 microns, though some can vary by two orders of magnitude and have variable width to height ratios. In stock configurations, both these cytometers use a sensitive PMT to detect SSC, and therefore a minimum SSC threshold was used to fully capture a wide range of particle sizes.

Bacteria must then be distinguished from small debris particles in the buffers and the sample itself, particularly in faecal samples which contain substantial autofluorescent debris. Cell-permeable SYBR Green I nucleic acid stain identified bacteria in samples from specific pathogen-free (SPF) mice but did not stain filter-sterilized FACS buffer (Fig. S1A). Nucleic acid staining may be variable amongst diverse bacteria. This is because there may be differences in cell permeability and transport mechanisms, the size and/or number of copies of genomic DNA, and any individual cell may be undergoing asymmetric division. These could produce a wide distribution of SYBR staining intensities. To ensure these are captured whilst minimising contaminating debris, stool from germ-free (GF) mice and unstained SPF stool was used to define SYBR+ gating parameters specific for bacteria (Fig. S1A).

Next, we sought to validate the specificity of the PE-conjugated rat anti-mouse IgA (mA-6E1) antibody that would be used in downstream IgA-Seq experiments. To do this we used IgA deficient mice as negative controls and a strain specific IgA antibody as a positive control. SPF *Rag1*^−/−^ mice have a complex microbiota but lack T cells and mature B cells responsible for the production of IgA. Essentially no bacteria were stained by the PE anti-mouse IgA antibody in samples from *Rag1*^−/−^ mice compared to roughly 4% of the microbiota in wild-type SPF mice (Fig. S1B). To further confirm the specificity of the IgA antibody, we replicated the approach of Kau et al. [18], using an *in vitro* system with an artificial 2-member bacterial community in which only one strain should be stained by PE anti-mouse IgA. *Bacteroides thetaiotaomicron* (*B. theta*), a common commensal of the murine gut, was coated with a mouse IgA antibody specific to the *B. theta* strain VPI-5482 [23]. *Escherichia coli* was coated with a mouse IgG antibody as a control and mixed with the labelled *B. theta* prior to staining with the PE anti-mouse IgA antibody. Flow cytometry revealed that a proportion equivalent to the expected *B. theta* population was stained by the PE anti-mouse IgA antibody (Fig. S1C), confirming the specificity of the antibody to be used in downstream IgA-Seq experiments. The gating strategy used for the IgA-Seq protocol (Fig. 1A) was validated using these SYBR and IgA staining controls.

**Figure 1.**
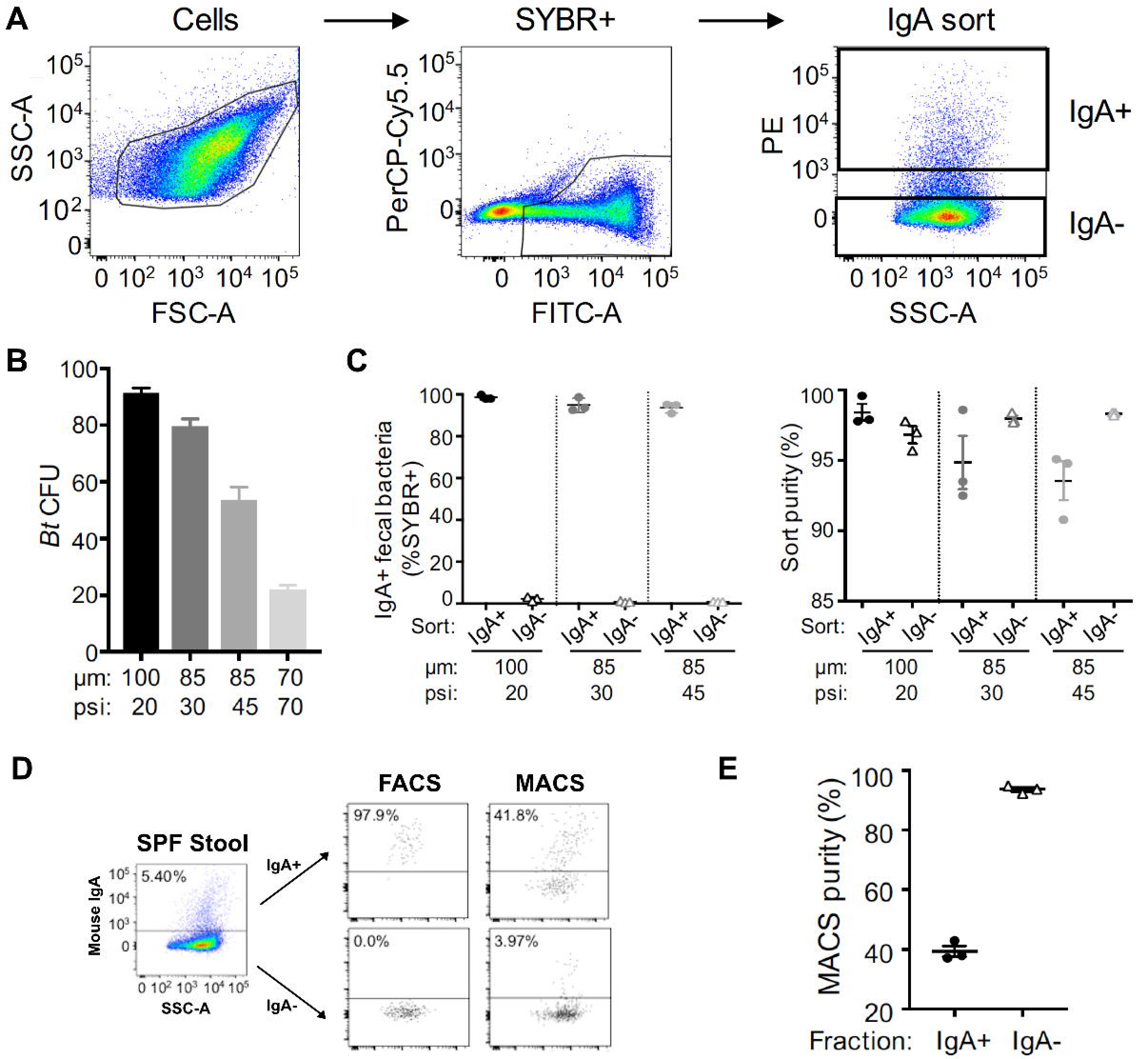
Benchmarking a protocol to isolate IgA+ and IgA- faecal bacteria by fluorescence activated cell sorting. A) IgA-Seq gating strategy. Faecal bacteria from SPF C57BL/6 mice stained with anti-mouse IgA (PE). Gating on a population based on FSC and SSC characteristics, followed by discrimination of SYBR+ (FITC) bacteria from autofluorescent (PerCP Cy5.5) debris. IgA+ and IgA- populations were sorted from SYBR+ cells. B) Viability of B. theta (*Bt*) collected by various nozzle sizes (μm) and sheath pressures (psi) on the FACSAria III. Culture-grown *Bt* was stained with mouse anti-*Bt* IgA (clone 255.4) and rat anti-mouse IgA (PE, clone mA-6E1). 100 cells were plated in Single Cell precision mode onto BHIS agar plates for enumeration of colony forming units (CFU). C) IgA+ and IgA- fractions were collected from SPF C57BL/6J mice (n=3). *Left:* percentage of IgA+ faecal bacteria detected in each collected fraction for a range of nozzle and pressure configurations. *Right:* sort purities for these configurations defined by the percentage of events that remained in the designated sort gates. Lines represent means with SEM. D) A portion of the IgA-stained faecal suspensions from Fig. 2A were also separated into IgA+ and IgA- fractions by magnetic activated cell separation (MACS) as a comparison. PE IgA- stained samples were incubated with anti-PE microbeads and fractions collected by LS column separation. Representative flow cytometry plots of the starting material in SPF stool and the IgA+ and IgA- fraction isolated by FACS and MACS. E) MACS purity of collected fractions as determined by flow cytometry, technical triplicates from the same stool suspension.

### A benchmarked protocol for IgA-Seq

The goal of IgA-Seq is to isolate bacterial populations coated by endogenous host IgA and to identify them by 16S rRNA gene sequencing. To this end, it is critical to identify FACS parameters that maximise the integrity and purity of sorted samples. Using *B. theta* as a model commensal anaerobe, a culture-grown suspension of *B. theta* was once again labelled with strain-specific IgA and PE anti-mouse IgA, and 100 cells were sorted as an array onto an agar plate to determine viable colony forming units (CFU). The default cytometer settings for 100 and 85 micron nozzles were tested, in addition to a custom 30 psi configuration for the 85 micron. Intact bacterial recovery decreased with smaller nozzles sizes and higher pressures, and viability was enhanced by sorting at the lower 30 psi pressure on the 85 micron nozzle (Fig. 1B). Only 20% of bacteria could be recovered with the 70 micron nozzle (Fig. 1B), which was discontinued from further testing.

To test the sort fidelity of the fractions produced by each of the configurations, we sorted IgA+ and IgA- bacterial populations from SPF mouse stool (Fig. 1A). These fractions were then analysed using the same parameters to determine the percentage of bacteria that were retained within defined sort gates. All three FACS configurations enabled >90% sort purity for both IgA+ and IgA- fractions (Fig. 1C). This is in contrast to magnetic activated cell separation (MACS), an alternative sorting modality which performed poorly with purities of 37.1– 42.9% for the IgA+ fraction (Fig. 1D-E). The FACS configuration utilising the 100 micron nozzle produced the highest sort purities in both IgA+ and IgA- fractions, followed by the 85 micron nozzle at 30 psi (Fig. 1C). Practically, the low pressure required for the 100 micron nozzle reduces the flow rate and throughput by ∼70%. Given that a minimum of 400,000 cells are required per fraction for downstream sequencing applications, the 85 micron with 30 psi configuration is vastly more time effective while maintaining high viability and sort purities. We also found this configuration allowed for sensitive sort control that, if desired, allows for finer grain collection of IgA+ subpopulations for instance IgA^bright^ and IgA^dim^ fractions (Fig. S2). This configuration was selected for downstream IgA-Seq.

As a final test of the precision of our sorting approach, we devised an experiment to specifically isolate an IgA+ strain from a complex stool microbiota. We once again took advantage of our ability to coat *B. theta* with mouse IgA in a strain-specific manner, which we stained separately before spiking into an unstained faecal suspension prepared from SPF wild-type mice (Fig. 2A). Endogenous IgA-coating of faecal microbiota remained but as the faecal suspension was not stained only the PE anti-mouse IgA stained *B. theta* culture should be isolated with this approach. *B. theta* comprised roughly 12.5% of the whole community in the spiked stool mixture, and high sort purities were achieved for both the IgA+ (97.4% – 99.4%) and IgA- fractions (97.3% – 100%) across three replicate sorts Figure 2B). 16S rRNA gene sequencing was performed on the complex stool community prior to the addition of *B. theta*, and in addition to the IgA+ fractions after three sorts from the same spiked sample. Genomic DNA from *B. theta* was included as a positive control. As a contamination check, sheath fluid was also included but it contained little DNA and did not produce a viable library. The IgA+ fractions were comprised almost entirely of *B. theta* (98.4 – 99.6%; Figure 2C-D) and lacked any additional unstained taxa observed in the complex community. This demonstrates that our optimised protocol can sort IgA bound bacteria from a complex gut microbiome with high accuracy.

**Figure 2.**
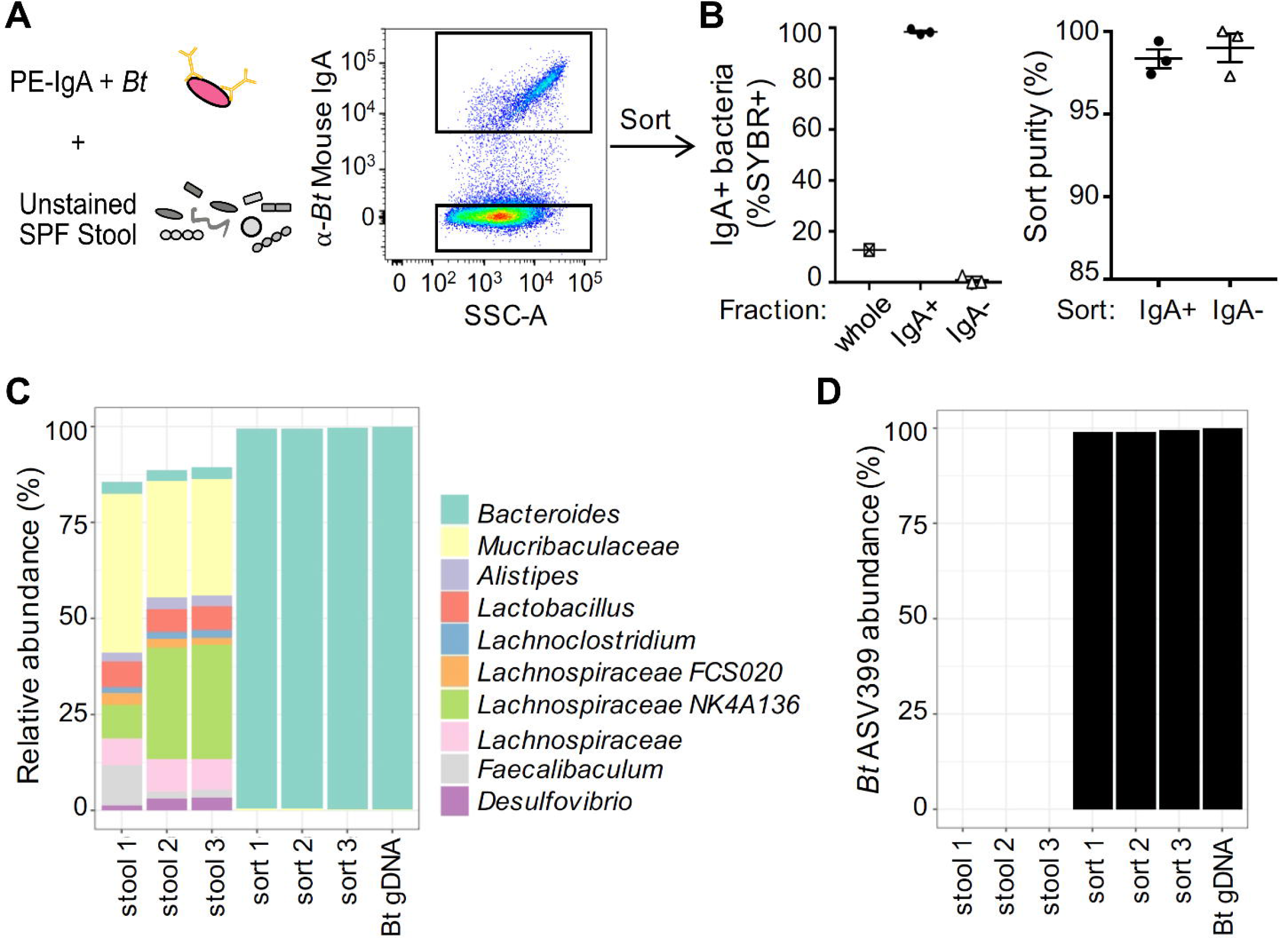
Testing IgA-Seq specificity in a complex faecal microbial community. Culture-grown *Bt* was stained with mouse anti-*Bt* IgA (clone 255.4) and rat anti-mouse IgA (PE, clone mA-6E1), then spiked into an unstained faecal suspension prepared from SPF C57BL/6J mice with a complex microbiota. A) Experimental design and sort setup, gated on SYBR+ cells. B) Sorts repeated on the same *Bt*-stool suspension to generate technical triplicates. *Left:* percentage of IgA+ bacteria in the pre-sorted sample, IgA+ and IgA- fractions. *Right:* purity check on the collected fractions. Lines represent means with SEM. C) Taxa abundances based on 16S rRNA gene sequencing of a complex stool microbiome (triplicate sequencing replications stool 1-3), triplicate sorted IgA+ fractions (sort 1-3), and *Bt* genomic DNA (*Bt* gDNA). Top 10 genera are shown. D) Relative abundance of *Bt* ASV399 in sorted samples.

### Pre-sort taxonomic abundances influence existing IgA binding scores

Once a sample has been sorted, amplicon-based sequencing of extracted DNA from the sort fractions is used to determine the relative abundances of the taxa within them. As discussed previously (see Background), the abundance of a taxon in the IgA+ fraction is a function of both its likelihood of IgA binding and its overall abundance in the pre-sort community. As we are principally interested in the former, we must use scoring metrics to quantify the relative IgA binding of taxa whilst taking their pre-sort abundances into account. To date, this has principally been achieved using indices that compare taxon abundances in the IgA+ and IgA- fractions and two such indices have been described – the Palm and Kau indices (see Methods). Using a toy example where two samples have identical IgA binding affinities for two species but differ in their starting abundances, we can see that these scores are still influenced by pre-sort abundances and produce different affinity scores for each sample (Fig. 3A). This is because these scores do not account for the fact that abundances are relative measures and thus the starting abundance and IgA binding of all other taxa will also influence the observed abundance of a given taxon in the sorted fractions.

**Figure 3.**
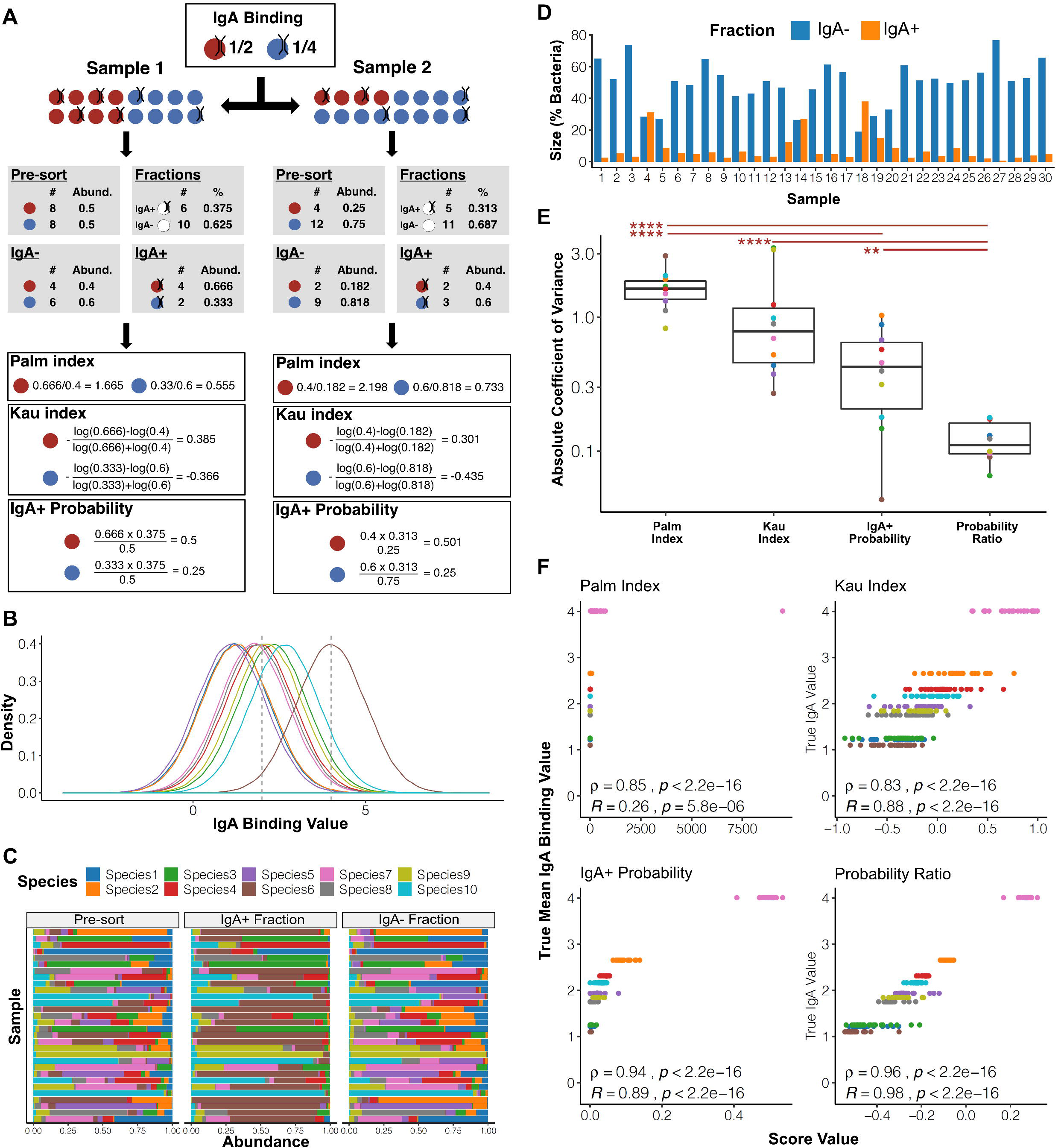
The influence of pre-sort abundances on existing IgA-Seq scores can be overcome using probability-based approaches. A) A toy example where two samples have two species with the same IgA binding profiles but different starting abundances. B) The distribution of arbitrary IgA binding values used for the ten species in the IgA-Seq simulations, dashed lines indicate the IgA+ and IgA- fraction thresholds. Species coloured as in C. C) The species abundances in the pre-sorted, IgA+ and IgA- samples for the 30 samples simulated. D) The size of the IgA+ and IgA- fractions in the simulated samples as a percentage of their total bacteria. E) Comparison of variance in IgA binding estimates across samples using each scoring approach. Coefficient of variance was calculated for each species individually. ** = p<0.01, **** = p<0.0001 in Mann-Whitney tests after FDR correction. Boxes represent the median and interquartile range (IQR) and the whiskers the largest and smallest values within one and half IQR of the upper and lower IQR limits. F) Pearson and Spearman correlations between the true mean IgA binding scores used to simulate data and the scores estimated by each of the different indices.

If we instead consider the IgA+ relative abundance as the probability of a taxon being within the IgA+ fraction, it is possible to apply Bayes’ theorem to derive a posterior probability of IgA binding for each taxon, i.e. the probability that a given bacterium will be bound to IgA given that it belongs to a given taxon (see Methods for a more detailed discussion). In our example, this posterior probability is not influenced by the starting abundances and provides a direct quantification of the likelihood of IgA binding for each taxon (Fig. 3A). Here, we describe two scoring methods based on these posterior probabilities: 1) The IgA+ probability – using the positive fraction probability directly and 2) The Probability Ratio – the ratio of the IgA+ and IgA- probabilities (see Methods).

### Probability-based scores accurately recapitulate relative IgA binding in simulated data

As the true level of IgA binding to each bacterial taxon cannot be measured directly *in vivo*, we created a platform to simulate an IgA-Seq experiment *in silico* to benchmark scoring methods. This allowed IgA-Seq data to be generated with pre-defined relative levels of IgA binding between species. In each simulation we maintain the relative level of IgA binding for each species across all samples and only change the pre-sort abundances of species. This allows us to determine how robust scoring methods are to these effects. A perfect scoring method should produce identical IgA binding scores for each species across all samples, with a relative ranking of IgA binding between species that matches the pre-defined levels.

Ten species were simulated, each having a unique normal distribution of IgA binding. IgA binding was represented by an arbitrary value that just represents the relative levels of binding between taxa (Fig. 3B). 30 samples were generated each containing 100,000 bacteria. The bacteria were assigned to each of the ten species randomly to produce different log distributed starting abundances for each sample (Fig. 3C). Each bacterium in each sample was then assigned an IgA binding value by sampling from the distribution of its species. Bacteria whose binding values were above four were considered IgA+ and those below two in the IgA- fraction. These thresholds were chosen based on the overall distribution of IgA values to mimic the extremes selected in a typical IgA-Seq experiment (Fig. 3B). The number of bacteria in each sample’s IgA+ and IgA- fraction was used to determine the size of the fractions and the species counts were converted to taxonomic relative abundances (Fig. 3C). These values were in turn used to derive the various IgA binding scores.

Samples had different IgA+ and IgA- fraction sizes, defined as the percentage of total bacteria within the fraction, due to their different initial taxonomic compositions and the differing IgA binding of the ten species (Fig. 3D). Given that each taxon’s IgA binding was constant across samples, we would expect there to be no variation across samples if an IgA binding score properly adjusts for the differences in sample composition. Comparing the consistency of scoring methods across samples, the IgA+ Probability had a significantly lower coefficient of variation than the Palm index and the Probability Ratio had significantly lower variation than all other scores. The probability-based methods were more consistent than the Palm and Kau indices, and thus less influenced by the differences in pre-sort abundances between samples (Fig. 3E).

When considering the relative rankings of the species, all four scoring methods were significantly correlated with the true IgA binding affinities and ranked species in the correct order of IgA binding (Fig. 3F). However, the Palm and Kau indices had a lower correlation than either of the probability-based methods due to the larger variation between samples. Comparing the results of Pearson and Spearman correlations, it is apparent that the Kau and Probability Ratio scores produce a more linear distribution than the other two methods. This is due to their consideration of both the IgA+ and IgA- fractions, which is not in the case in the IgA+ Probability, and taking the log to centre the score, which is not the case in the Palm index or IgA+ Probability. Overall in this initial simulation, probability-based methods proved robust to the influences of pre-sort abundances, which introduced variability to the Palm and Kau indices. The Probability-Ratio almost perfectly recapitulated the relative relationships of the true IgA binding values.

### Probability-based methods overcome biases effecting the interpretation of between group comparisons

The Kau and Palm indices showed variation in scores due to differences in pre-sort abundances. This could be particularly problematic when comparing IgA binding between groups that have inherent and consistent differences in microbiome composition. This is likely, as a key use-case for IgA-Seq is to determine if previously established microbiome differences between groups, such as healthy controls and disease patients, are being driven by differential targeting of the IgA pool.

To determine the effects of pre-sort abundance biases on inter-group comparisons, we repeated the simulation but divided samples into a case and control group. The simulation was run with 30 samples per group each containing ten species with the same underlying IgA affinities as before (Fig. 3B).

Random initial abundances were then generated as previously but additional Species 10 (with a moderate IgA affinity value) was introduced to the 30 case samples – simulating outgrowth of one specific taxon in a disease. In this simulation, as before, an optimal scoring approach will not detect any differences between the cases and controls whose species share the same IgA binding and only differ in their pre-sort abundances.

As expected, Species 10 constituted a larger percentage of the relative abundances in both the IgA+ and IgA- fractions of the cases compared to the controls (Fig. 4A). This led to a significantly higher IgA- fraction size in the control group when compared to cases but no difference in the IgA+ fraction sizes (Fig. 4B). Comparing the overall distribution of scores, there was greater variability across samples in the Kau index than the Probability Ratio, and Species 10 IgA binding estimates were higher in cases using the Kau index (Fig. 4C). When testing differences between the simulated case and control groups, the Kau index inferred significant differences in the IgA binding of both Species 10 and Species 6 where there should be none (Fig. 4D). This is an artefact of measuring relative abundances. In the simulated disease cases there was an increased abundance of Species 10 in the IgA+ fraction. This reduces the relative contribution of Species 6 (the highest affinity taxa that would otherwise constitute the majority of the IgA+ fraction) and alters the ratio when calculating the Kau score. No significant differences were observed using the Palm index but this only resolved the highly bound Species 6, which displayed increased variance in the controls (Fig. S3). The Probability Ratio resolved both high and low IgA binding of taxa and was more resistant to the influence of the pre-sort abundances, maintaining similar estimates between cases and controls (Fig. 4D).

**Figure 4.**
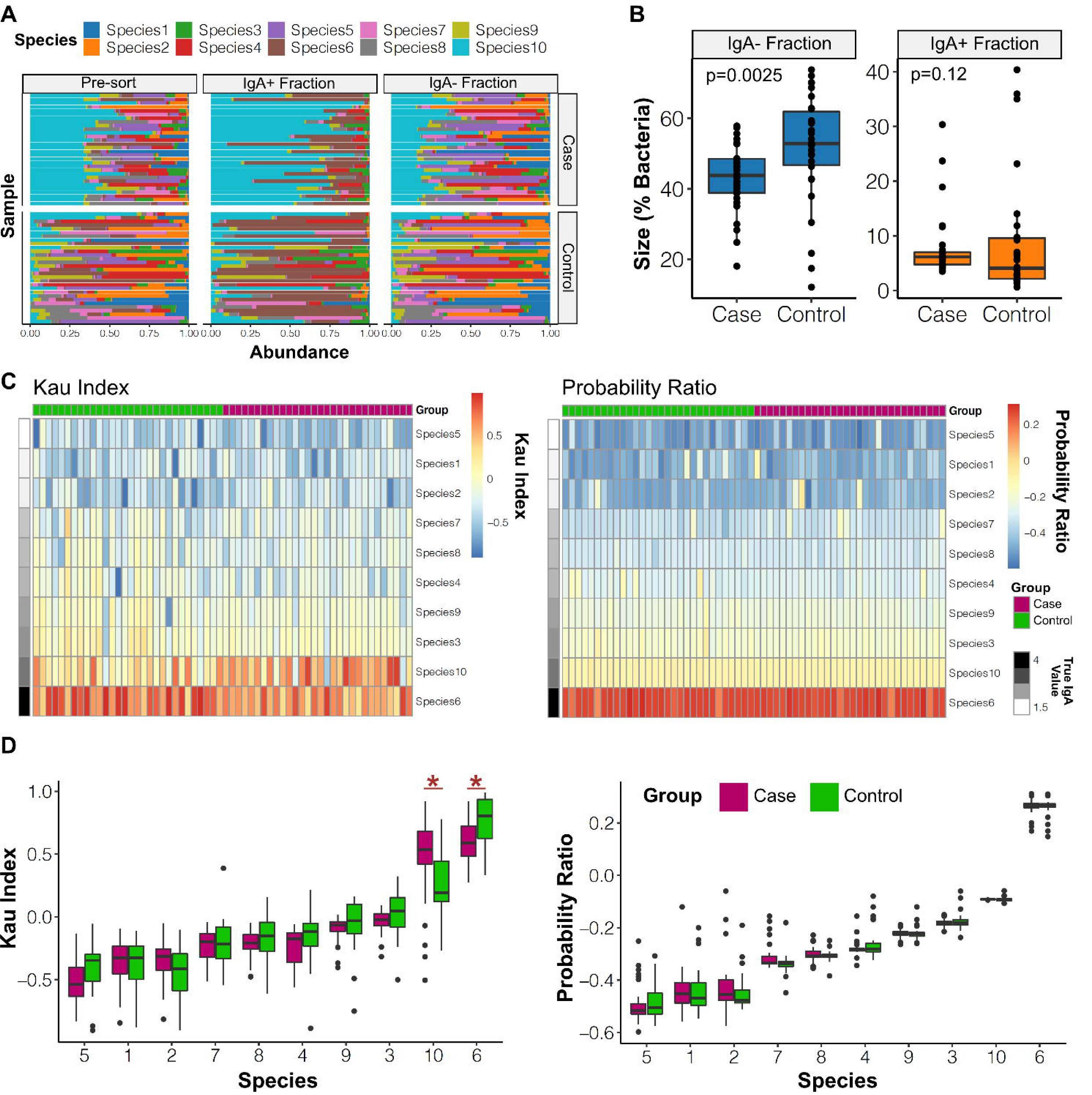
The probability ratio overcomes pre-sort abundance biases when carrying out between-group comparisons. A) Simulations were carried out as in Figure 4 (Controls) with an additional 30 samples that were initiated with the additional abundance of Species 10 (Cases). Abundances in the pre-sort, IgA+, and IgA- fractions using the thresholds as in Figure 4A. B) Differences in the IgA+ and IgA- fraction sizes as a percentage of total bacteria between cases and controls. P-value shown from Mann-Whitney tests. C) Heatmaps showing the scores estimated by the Kau and Probability Ratios across all samples. D) Comparison of IgA binding scores for the ten species in cases and controls when using either the Kau score or the Probability Ratio. Significant differences are highlighted with * (FDR adjusted Mann-Whitney p<0.05) and species are ordered by true IgA binding value. In all boxplots the boxes represent the median and interquartile range and the whiskers the largest and smallest values within one and half IQR of the upper and lower IQR limits. In D, points outside this range are shown.

### IgA-Seq in a mouse model of colitis produces different outcomes across scoring methods

To demonstrate the utility of our benchmarked IgA-Seq protocol we applied it to study commensal IgA binding in a mouse model of colitis. *Helicobacter hepaticus* (*Hh*) can inhabit the murine intestinal tract without harm but can emerge as a pathogen when host immune responses are dysregulated as a result of either genetic susceptibility or targeted immunomodulation [24]. Abrogation of anti-inflammatory IL-10 signalling by administration of an anti-IL10 receptor antibody (aIL10R) results in *Hh-*driven colitis[25]. We recapitulated this model in a gnotobiotic C57BL/6 mouse colony stably colonized with a 12-member microbiota (MM12) that recapitulates the taxonomic diversity of a more complex community[26]. This provides a defined minimal microbiota to specifically interrogate IgA binding.

Control MM12 mice treated solely with weekly aIL10R injections did not develop any pathology (Fig. 5A). In contrast, MM12 mice that received aIL10R in conjunction with *H. hepaticus* infection (*Hh*+aIL10R) developed marked inflammation and colitis after 14 days (Fig. 5A). Lipocalin-2, released by activated neutrophils and a biomarker for inflammation[27], was also elevated in the colon contents of *Hh*+aIL10Rmice but not control aIL10R only mice, whose lipocalin-2 levels were similar to steady-state mice lacking aIL10R injection (Fig. 5B). There was a significant increase in IgA-coating of colonic bacteria from colitic mice (Fig. 5C-D). Colonic contents from both control and colitis groups were subjected to IgA-Seq to identify taxa targeted during inflammation.

**Figure 5.**
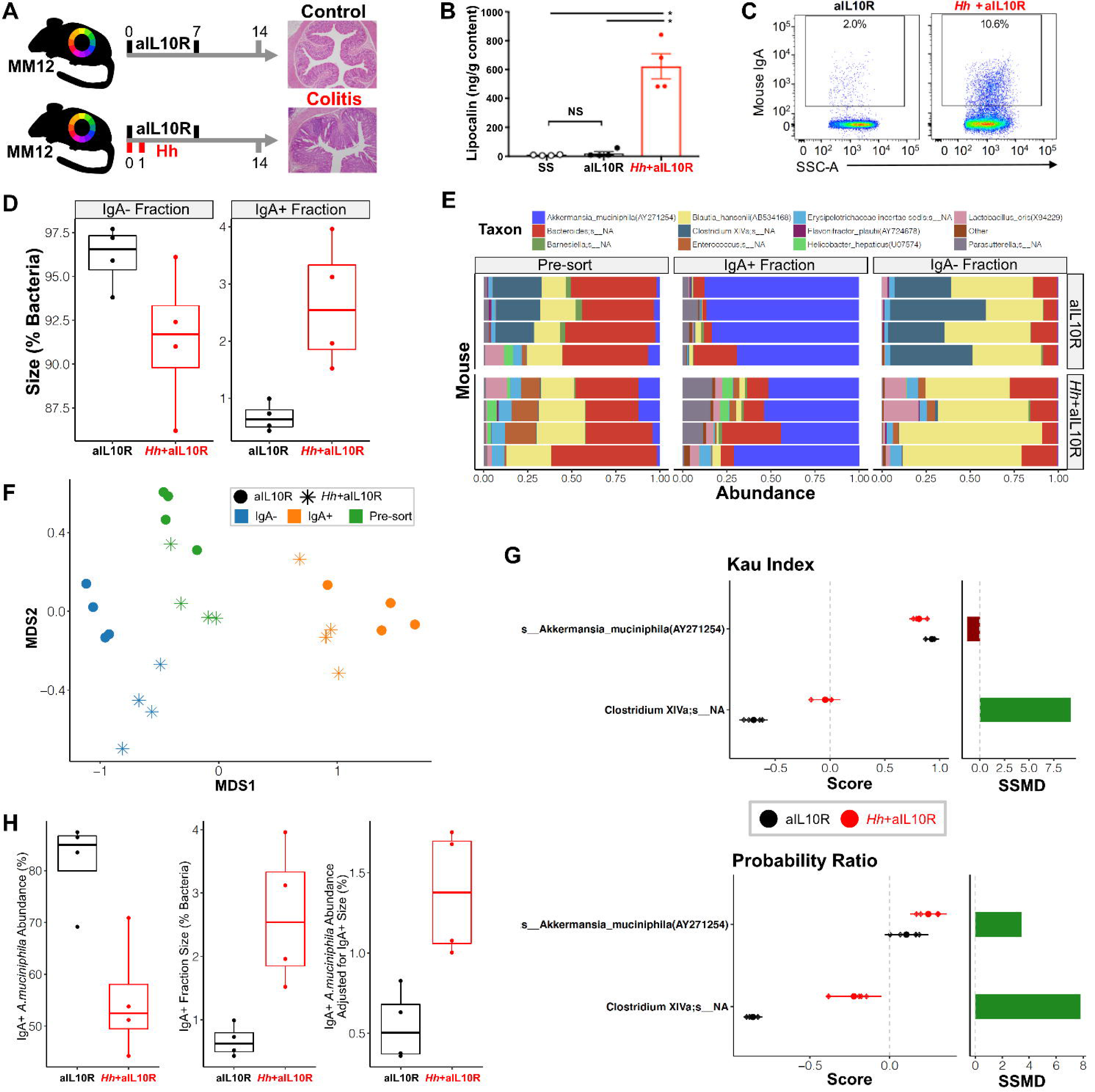
IgA-Seq in a mouse model of colitis with a defined 12-member microbiota. MM12 mice were administered weekly injections of anti-IL10R (day 0, 7). Controls received anti-1L10R alone (aIL10R, n=4), and the colitis group were additionally infected with *H. hepaticus* at the start of the experiment (Hh+aIL10R, n=4). A) Experimental design and representative colon histology with haematoxylin and eosin staining. B) Mouse lipocalin-2 ELISA as a measure of colonic inflammation. Lines at means and SEM. **p<0.01 unpaired Mann-Whitney test, ns= non-significant. SS represents steady state mice with no aIL10R exposure. C) Representative flow cytometry plots of the percentage of colonic MM12 bacteria coated by IgA. D) Percentage of bacteria coated by host IgA and subjected to IgA-Seq. P-values represent Mann-Whitney tests. E) Relative abundance plots showing the MM12 species in each sample. Each row represents an individual animal. F) Non-metric multi-dimensional scaling of pre-sort, IgA+, and IgA- fractions from Bray-Curtis distances. G) *Left:* means and 95% confidence intervals of IgA binding scores significantly different between groups. Significance determined by complete permutation of group labels where, exact permuted p<0.1). *Right:* Mean difference of scores between groups (colitic – control). Mean is normalised by its standard deviation for comparability between scoring methods (using the strictly standardised mean difference SSMD). H) *Left*: boxplots of *A. muciniphila* relative abundance. *Centre*: boxplots of IgA+ fraction size. *Right:* boxplots of adjustment for the difference in fraction size (abundance in the IgA+ fraction size multiplied by *A. muciniphila* abundance). In the boxplots in D and H, boxes represent the median and IQR and the whiskers the largest and smallest values within one and half IQR of the upper and lower IQR limits.

Taxa were profiled in the pre-sort samples and the IgA+ and IgA- fractions using 16S rRNA gene sequencing. Analyses were carried out at the level of species summarised amplicon sequence variants (ASVs). Using the V4 16S rRNA gene region we were able to distinguish almost all members of the MM12 and *Hh* either by species level assignment or sole membership of a genus (Table S1). The exception were two closely related *Clostridium* species that were both assigned to *Clostridium XlVa*. A blank sequencing control was used to identify potential contaminants introduced at the DNA extraction and library preparation stage. Given that we used mice with a defined microbiota, we were also able to determine the presence of contaminants introduced elsewhere. After removing taxa observed in the blank control, several unexpected (non-MM12 or *Hh*) species remained (Fig. S4A). The majority of these were at a lower abundance than the expected species across all samples. These were removed from later analyses using an abundance threshold chosen to remove rare taxa whilst retaining the majority of total observations (Fig. S4B). It is of note that contaminant taxa were far more predominant in the sorted IgA+ and IgA- fractions. This may be due to the lower biomass in these samples but could also be due to species being introduced during sorting or library preparation. These were accounted for by additional screening to only retain species observed in the pre-sort sample for each mouse, which led to no unexpected taxa being observed in any sample (Fig. S4C). Thus, we recommend using abundance thresholding followed by validation against pre-sort samples to filter taxa prior to scoring in IgA-Seq analyses.

After quality control, *Hh* was detected in the whole and IgA+ fractions of all four *Hh+*aIL10R mice and samples clustered clearly by both experimental condition and sort fraction (Fig. 5E-F). *Hh* was also detected in the pre-sort sample in one of the aIL10R only mice, but as this was not detected in either the IgA+ or IgA- fractions it ultimately did not influence the IgA binding scores for this mouse. We compared the different IgA binding scores across all mice, focusing on the methods that included both the IgA- and IgA+ fractions. All methods broadly ranked species IgA binding in the same order and there was a significant positive correlation between all scores (Fig. S5A). The Kau index and Probability Ratio shared the strongest correlation, with the main difference between the two being a more even distribution of values within the Probability Ratio than in the Kau index, which contained isolated peaks of scores where taxa were predominantly observed in the IgA+ or IgA- fraction (Fig. S5A).

When comparing species IgA binding scores between groups, the method used had important and distinct effects on the interpretation of the results. All methods identified significant differences in IgA binding of *Clostridium* between the *Hh*+aIL10R and aIL10R only treatments (Fig. 5G, Fig. S5B). This was the largest effect observed and was clearly detected by all three methods in the same direction with an increased IgA binding of *Clostridium* in colitic mice. Some taxa, such as *Enterococcus*, were only observed in the IgA+ and/or IgA- fractions in the colitic mice. It was therefore not possible to compare their binding relative to the control mice. This highlights the need to additionally observe presence/absence patterns of taxa in the fractions alongside using quantitative scoring methods.

The Kau index and Probability Ratio additionally detected a significant difference in IgA binding to *A*.*muciniphila* between the groups. However, the Kau and Palm scores were lower for *A*.*muciniphila* in colitic mice, whereas the Probability Ratio was higher (Fig. 5G). This discrepancy results from the large contribution of *Hh* to the relative abundances in the IgA+ fraction of the colitis group. This inherently reduced the observed relative abundance of *A. muciniphila* in this fraction even in the absence of a change in the absolute levels bound to IgA. This is less pronounced in the IgA- fraction (where less *Hh* is observed) and thus the overall IgA+/IgA- ratio of *A. muciniphila* was reduced (Fig. 5G). However, even though *A. muciniphila* accounts for a reduced percentage of the IgA+ fraction, the total IgA+ fraction is much larger in the colitic mice (Fig. 5H). A smaller percentage of this much larger total IgA+ fraction actually represents a larger total amount of *A. muciniphila* bound to IgA in the colitic group (Fig. 5H). Thus the Probability Ratio, which accounts for the fraction sizes, infers a positive rather than negative shift in IgA binding of *A*.*muciniphila*. This highlights the benefit of using probability-based metrics that provide a direct estimate of the likelihood of IgA binding for each taxon in comparison to solely abundance-based indices, which might lead to a conflicting interpretation without careful consideration of what the scores represent.

### The Probability Ratio increases power to detect differences in IgA binding between conditions

As an additional test of the Probability Ratio we applied it to a published IgA-Seq dataset with extensive experimental evidence to support differences in IgA binding between conditions. Huus et al. used FACS-based IgA-Seq to study bacterial binding of GI IgA in a mouse model of nutrient restriction [20]. Mice on a conventional (CON) or moderately malnourished diet (MAL) were tracked from weaning to adulthood. Applying the Kau index, they found that mice on a conventional diet develop increased binding of *Lactobacillus* as they age from three to seven weeks old, whereas malnourished mice do not. They observed this effect to be consistent across different sites in the GI tract and confirmed it was driven by differential binding of IgA to *Lactobacillus* using ELISA. This provided an ideal dataset to test the robustness of the Probability Ratio.

Huus et al. provided processed 16S rRNA gene sequencing data generated from faecal samples used in their study and the associated IgA+ and IgA- fraction sizes, which had been recorded for a subset of the original data [20]. This enabled the Probability Ratio to be calculated for samples from four CON and MAL mice at each of the three ages. The Kau score (used in the original study) was also calculated as a comparator.

Comparing between diet and ages using the Kau index, we identified five taxa with differential IgA binding (Fig. 6A). *Lactobacillus* was the only taxon significantly different by the interaction of age and diet and reflected the patterns observed previously. Three taxa, *Lactococcus* and unclassified *Lachnospiraceae and Erysipelotrichaceae*, displayed increased IgA binding in MAL mice. All three of these taxa were observed with the same association in the original analysis of the dataset. We additionally observed a reduced IgA binding of *Roseburia* at three week old MAL mice. This was not reported in the Huus et al. analysis using the Kau index. However, due to the smaller sample size available we explicitly targeted our testing to between group effects and the prior analysis did not.

**Figure 6.**
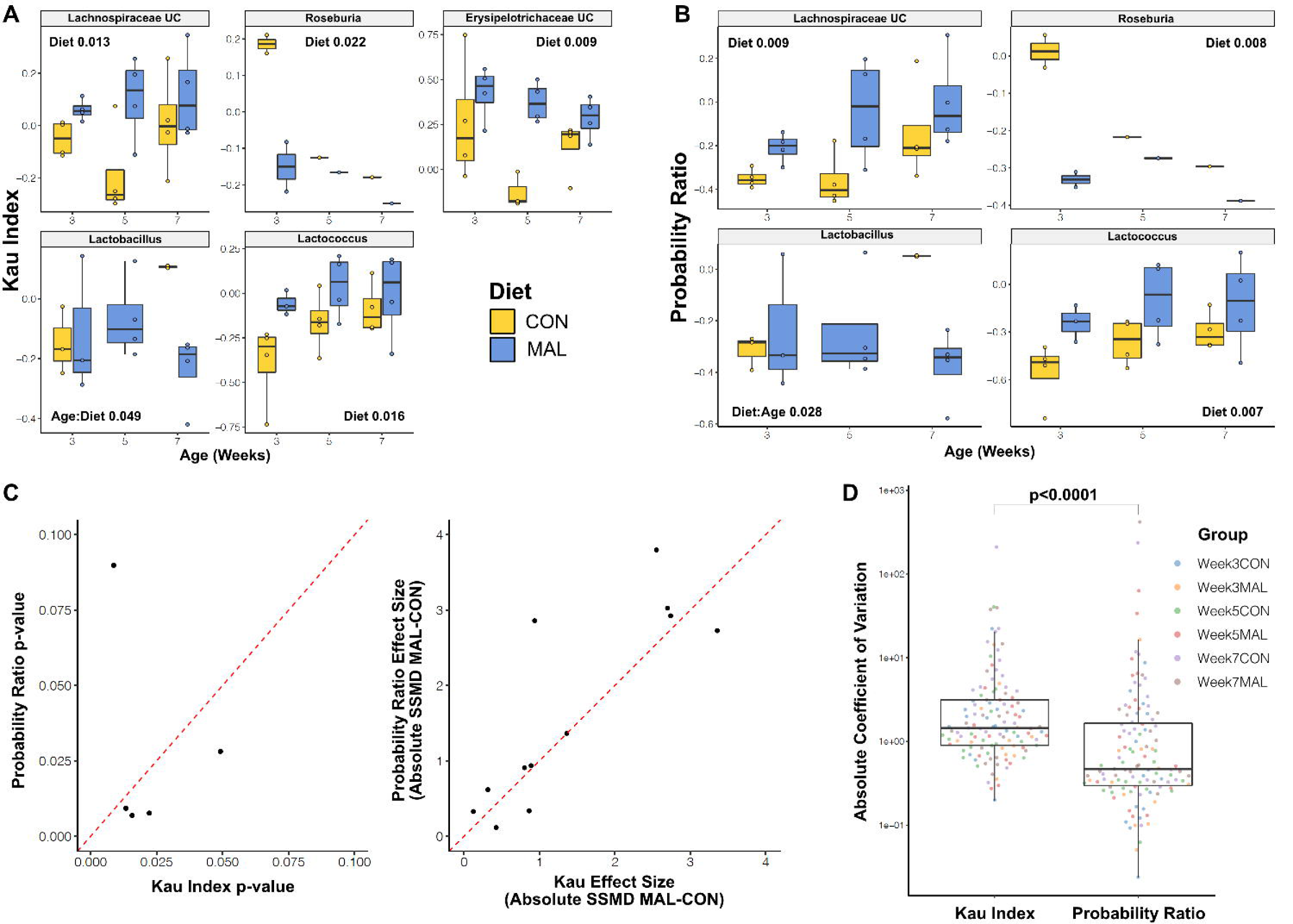
Increased power when using the Probability Ratio to reanalyse data from Huus et al. 2020. A) Boxplots showing the Kau index scores at each age for all taxa that were significantly different by either diet or the interaction of age and diet in two-way ANOVAs (nominal p<0.05). Each plot shows which of diet and the diet:age interaction were significant and their p-values. Boxes represent the median and IQR and the whiskers the largest and smallest values within one and half IQR of the upper and lower IQR limits. UC represents taxa that were unclassified below the given taxonomy. B) Plot as in A but for the Probability Ratio scores. C) Left: Scatterplot of the p-values for all the comparisons that were significant in either A or B showing the p-value when using the Probability Ratio (y) or Kau Index (x). The red-line shows y=x, highlighting the lower p-values in the Probability Ratio tests. Right: Scatterplot as for p-values but showing the effect size (difference in means between the two diets) at each of the three ages for all of the significant comparisons in A and B. Effect size is quantified as the strictly standardised mean difference (MAL-CON). D) Boxplots comparing the absolute coefficient of variation of each taxon’s score within each experimental group (for all taxa detected in the Huus et al. dataset) when using either the Kau Index or Probability Ratio. Significance shown from Mann-Whitney U test.

Carrying out similar modelling using the Probability Ratio to score IgA binding, we reproduced all of the associations identified using the Kau index with the exception of the unclassified *Erysipelotrichaceae* taxon, which was no longer significantly different between groups (Fig. 6B). Across all of the other associations detected as significant by either the Kau Index or Probability Ratio, the Probability Ratio consistently produced lower p-values (Fig. 6C, left). This reflects an increased magnitude in the differences between the mean scores of the MAL and CON groups in most comparisons when using the Probability Ratio compared to the Kau index (Fig. 6C, right). This can in part be attributed to a reduced variation in taxon scores within groups when using the Probability Ratio (Fig. 6D), as was also observed in the simulated data (Fig. 3E). This increased power is an additional benefit of accounting for the size of the IgA+ and IgA- fractions when scoring IgA-Seq data.

## Discussion

The development of IgA-Seq has enabled powerful, unbiased investigation of immune interactions between host and commensal microbiota. As this technique gains wider adoption, it is critical to benchmark experimental and analytical approaches to ensure accurate quantification and interpretation of IgA targeting. To this end, we provide an optimised IgA-Seq protocol and novel scoring approaches that facilitate accurate identification of IgA coated bacteria. We developed a simulation platform that enabled the first formal testing of IgA-Seq scores. This revealed that existing scoring approaches are influenced by relative abundances and, without careful consideration, can lead to the inference of alternate IgA binding patterns solely as a result of pre-sort abundance shifts. Instead, we have developed new probability-based scoring methods that overcome these limitations by directly quantifying of the likelihood of a taxon being bound by IgA. Importantly, as we have showed in both simulated and *in vivo* datasets, these probability-based scores provide increased power to detect differences in IgA binding between groups and promote accurate interpretation of IgA targeting of taxa in health and disease states.

Analysis of *in silico* and *in vivo* datasets revealed that it is critical to properly account for both the pre-sort taxonomic composition of a sample and the relative nature of fraction abundances when comparing IgA targeting between case and control groups. Scoring indices that neglect to do so can infer artificial and/or conflicting shifts in IgA binding estimates between experimental groups. Simulating consistent group-wise shifts in pre-sort abundances resulted in significant differences in taxon level binding estimates when using the Kau index. It is not that the Kau scores are incorrect, but rather that they represent an abstract combination of both the IgA binding and abundance of the taxon scored, and the IgA binding and abundance of all other taxa in the sample. This inherently complicates their interpretation. In contrast, the probability-based scores adjust the taxonomic abundances by the number of bacteria in IgA+ and IgA- fractions. This overcomes the relative nature of quantification, which removes the influence of other taxa, and allows for proper adjustment for pre-sort composition. As a result, probability-based scores provide a direct measure of the likelihood that a bacterium is bound to IgA given that it belongs to a specific taxon. As shown with the increased binding of *A*.*muciniphila* in the colitis model, this simplifies the interpretation of between group changes in IgA binding when using the Probability Ratio.

Aside from artefacts that arise in between-group comparisons, all scoring indices were correlated when considering global patterns across samples. This suggests that taxa ranked as generally highly bound in previous studies are unlikely to change if quantified using the novel probability-based scores. However, we found that the influence of pre-sort taxonomic composition meant there was higher variability between samples when using the Kau and Palm scores compared to the probability-based methods. This increased variance reduces the power to detect effects in experimental data and resulted in a moderate increase in power when applying the Probability Ratio to the Huus et al. data. Maximising power by using probability-based metrics is particularly desirable given the high investment of experimental time required to generate IgA-Seq data, which can introduce a practical limit on sample sizes.

Our work highlights the importance of selecting scores based on the underlying biological question. If the primary interest is to simply identify which taxa have the most bacteria bound by IgA, then the IgA+ abundance of a taxon can be used directly. If the purpose is to identify which taxa are being more/less preferentially bound over other taxa, which is arguably more relevant when studying IgA targeting, then probability-based scores that account for the relative nature of taxonomic abundances should be used (Fig. S6). To select between using the IgA+ Probability or Probability Ratio: if the sole interest is to identify taxa most likely to bind IgA then the IgA+ Probability is sufficient and benefits from its direct interpretability as a probability of IgA binding (ranging from 0-1); otherwise the Probability Ratio provides a more generally applicable score that also provides better resolution of relative binding between taxa with lower IgA binding. For example, detecting the loss of *Lactobacillus* binding in undernourished mice detected in the Huus et al study[20]. The Probability Ratio additionally benefits from being centred on zero, so a positive or negative score indicates if the majority of a taxon’s bacteria fall into the IgA+ or IgA- fraction respectively. Score choice might also be based on available sequencing data. The IgA+ Probability requires sequencing of the IgA+ fraction and the pre-sort sample and the Probability Ratio requires sequencing of the IgA+ and IgA- fraction. However, sequencing of both the IgA+ and IgA- fractions and the pre-sort sample has often been applied in previous studies [17, 18]. As we have shown here, this provides additional benefits in the detection of contaminants introduced by sorting and overall compositional differences between experimental groups.

We have also presented an optimised IgA-Seq protocol that enables precise separation of IgA+ and IgA- gut microbiota, and their cultivation to support subsequent interrogation in gnotobiotic disease models and/or strain phenotyping. Our FACS-based protocol outperformed MACS in sort purity and benefits from innate quantification of the IgA+ and IgA- fraction sizes required to calculate the Probability Ratio. We employed a protocol design strategy that combined individual approaches applied in previous studies including the controls used for validation of the staining and gating strategy [18, 28]. To minimize technical variation between experiments, regular instrument calibration alongside staining controls and gating templates should be employed. We utilised the same antibody lot for all experiments within this study, but staining controls and gating strategies need to be verified in instances of lot-to-lot variation. In addition, we developed a novel positive control approach to testing protocol accuracy by successfully isolating pure IgA-bound *B. theta* from a complex intestinal microbiota. While access to specific flow cytometers may vary, the concepts in experimental design presented herein can be applied to validate IgA-Seq protocols on different sort modalities.

Similar approaches to IgA-Seq have also been used to study other isotypes, such as IgM and IgG[22, 29–31], and immunoglobulins isolated from other sites, for example quantifying GI bacteria bound by serum IgA[32]. Expanding beyond IgA targeting of the gut microbiota, in future work, our protocol design strategy and scoring approaches could be applied to investigate other sample types and immunoglobulins. Further work could also be done to explore other variables influencing IgA binding scores. For example, amplicon-based taxonomic abundances are inherently influenced by PCR biases, differences in 16S rRNA gene copy number, and the taxonomic resolution of the variable region amplified[33–35]. However, these are more general limitations of 16S rRNA gene sequencing and their impact should be minimised assuming they occur uniformly in the IgA+ and IgA- fraction. Mock sorts have previously shown that the sorting process itself has little influence on community structure[17].

Technical variation in the isolation of IgA+ bacteria will also influence IgA binding estimates. For example, using a more sensitive anti-IgA antibody or setting wider IgA+ gating in FACS will increase the size of the IgA+ fraction and inherently increase the likelihood a taxon will be IgA+. This will not influence relative scoring between taxa within a sample but it is important to maintain these variables across samples in an experiment and/or consider randomisation of samples across technical batches to avoid confounding of experimental groups. For similar reasons, absolute score values from any method are unlikely to be analogous between experiments, however the relative rankings of taxa should remain comparable. IgA-Seq is additionally limited to scoring relative, overall, IgA binding between taxa. Understanding what drives the observed differences, such as alterations in the spatial distribution of bacteria or changes in the affinities of host IgA, requires additional targeted experiments, such as fluorescence *in situ* hybridisation and enzyme-linked immunosorbent assays[10, 20].

We have developed an optimised experimental protocol and novel scoring methodologies for IgA- Seq. These enable robust and accurate identification of commensal bacteria targeted by the host immune response. The IgAScores R package facilitates the application of these scores and we have presented a guiding framework for selecting appropriate metrics. Furthermore, its simulation platform will aid the future development of IgA-Seq analytics. This work enhances the accessibility and quality of IgA-Seq studies, which should consequently uncover novel mechanistic insights into host-microbiota interactions and the microbial drivers of health and disease.

## Conclusion

Existing methods used to score taxon-level IgA binding in IgA-Seq experiments are influenced by the initial taxonomic composition of samples and the relative nature of their quantification. This inflates the variability in these scores and can have important consequences in between-group comparisons, where scores might infer false or contrasting associations as a result of systematic differences in microbiome composition between groups. Probability-based IgA scores can be used that overcome this by providing a direct estimate of the likelihood a bacterium will be bound by IgA given its taxonomy. These scoring methods are available in the IgAScores R package. Combined with the practical approaches described here, these analytical developments enhance our ability to study taxon-level immunoglobin binding of commensal microbiota; by increasing the power to detect associations through reduced variability across samples and by providing an accurate, direct, quantification of taxon binding from IgA-Seq data.

## Methods

### Indices for scoring relative IgA binding of taxa in IgA-Seq experiments

#### Palm index

In Palm *et al*.’s initial description of IgA-Seq they simply used the ratio of a taxon’s IgA+ to IgA- abundance in a score termed the IgA coating index[17]. Here, for clarity, we refer to this as the Palm index (*Palm*). If *IgA*^+^ and *IgA*^−^ represent taxon abundances in the respective fractions, for taxon *i* in sample *j* the Palm index is defined as in Equation 1, where if 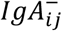 is equal to zero a pseudo count (*c*) is added to negate division by zero. Throughout we use a pseudo count of a similar magnitude but below the smallest abundance observed in any sample.

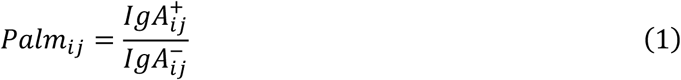

### Kau index

Kau *et al*. described the IgA index, which is also solely based on IgA+ and IgA- relative abundances [18]. It uses log transformation to make scores more comparable between taxa and considers the difference of the IgA+ and IgA- abundances relative to their total. This has the benefit of centering the score such that positive and negative values reflect overall higher abundances in the IgA+ or IgA- fractions respectively. Using notation as previously, the Kau index (*Kau*) is defined as in Equation 2.

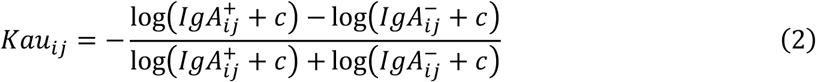

### IgA+ Probability

The relative abundances of taxa in a fraction or pre-sort sample sum to one. These can be considered as probabilities. For example, taxa abundances in the pre-sort sample represent the probability that any given bacterium in the sample belongs to a given taxon. The abundances in the IgA+ and IgA- fractions can be considered as conditional probabilities (i.e. the probability of event A given that B has occurred). For instance, IgA+ abundances represent the probability a bacterium belongs to a taxon given that it is sufficiently bound to IgA to be in the IgA+ gate. In the case of IgA-Seq we are interested in the inverse: the probability that a bacterium will be IgA+ if it belongs to a given taxon. This is a posterior probability and can be calculated following Bayes’ theorem (Equation 3).

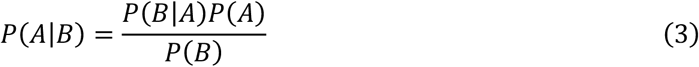

Where *P*(*A*|*B*) is the posterior probability that a bacteria is IgA+ given that it is from a chosen taxon. *P*(*A*) is the probability that any bacterium is bound to IgA sufficiently to be in the IgA+ fraction – measured as the percentage of bacteria that fall within IgA+ FACs gate. *P*(*B*) is the probability that a bacterium will belong to a given taxon – measured as the taxon’s relative abundance in the pre-sorted sample. *P*(*B*|*A*) is the probability a bacteria belongs to the taxon given that it is in the IgA+ fraction – measured by the taxon’s IgA+ abundance. In which case, the posterior probability 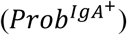 for taxon *i* in sample *j* is defined as in Equation 4.1. Where *FracSize* equals the fraction of sorted bacteria in the IgA+ gate and *PreSort* equals taxon abundances in the pre-sorted sample.

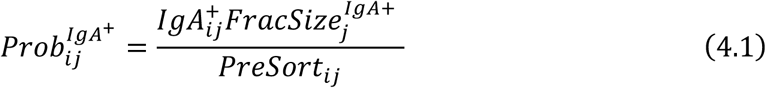

This measure will be referred to as the IgA+ Probability. The numerator provides a direct estimate of the proportion of initial bacteria bound to IgA and should theoretically always be smaller than the denominator (the total level of that taxon in the sample). However, due to amplification and other technical biases, this may not practically be the case in IgA-Seq data, thus Equation 4.2 can be used to account for this and ensure probability estimates fall below 1.

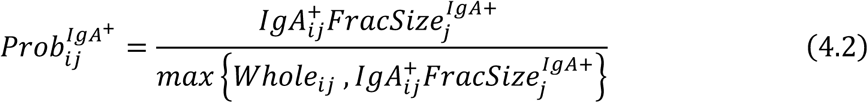

This will assign a probability of 1 when the estimate of IgA+ bacteria in a taxon is larger than the observed total abundance in the pre-sort sample, a reasonable assumption being that very high levels of IgA binding would be necessary to produce these results.

This score only requires two additional pieces of data than the Kau and Palm indices: the taxonomic abundances in the pre-sort fraction and the percentage of sorted bacteria that are in the IgA+ gate in the FACS run. The latter value is already generally recorded during sorting and most IgA-Seq studies to date have profiled the microbial communities from pre-sorted samples[17, 18]. Furthermore, if the only interest is to identify highly bound taxa this method can negate the need to sequence the IgA- fraction.

### Probability Ratio

The IgA+ Probability only provides a measure of the likelihood a taxon is sufficiently bound to IgA to be in the IgA+ fraction. The converse, the IgA- Probability, can also be calculated by simply substituting the IgA+ abundances and fraction sizes for the IgA- values. The IgA+ and IgA- probabilities can be combined to generate an IgA Probability Ratio. As both probabilities are normalised to the taxon’s abundance in the whole fraction it can be removed from the equation (Equation 5), negating the need to sequence the pre-sort sample if desired.

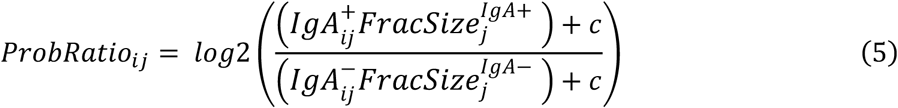

Pseudo counts (*c*) must be included in the Probability Ratio to negate division by zero where taxa are not observed in the IgA- fraction. Using the Probability Ratio has the benefit of capturing both extremes of IgA binding within a single score (whereas the IgA+ or IgA- Probabilities are limited to either direction) as well as, similar to the Kau index, centering values on zero by taking the log. This provides a clear indication of whether the majority of bacteria in a taxon are found in the IgA+ or IgA- fractions.

One limit of using the ratio as in Equation 5 is that the range of possible values returned is determined by the pseudo count used. The magnitude of these effects can be reduced by dividing *ProbRatio* by a scaler based on the pseudo count as in Equation 6. This will bring all values within the range −1 to 1 with these values representing the extreme case where all bacteria are in a single fraction and belong to a single taxon. Scaled Probability Ratios are used throughout.

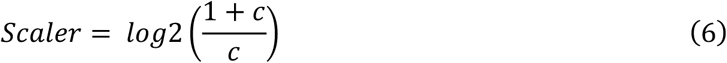

The Palm, Kau, IgA+ Probability, and Probability Ratio scores were all calculated as defined above using the *igascores* function that we have created as part of an R package IgAScores, which is available at *https://github.com/microbialman/IgAScores*.

### Simulating an IgA-Seq experiment

A simulation was used to generate IgA-Seq data where the underlying relative taxonomic binding of IgA binding was known. Ten species were assigned a mean IgA binding value by random sampling from an exponential distribution (raising two to the power of the generated values to increase their spread). These values are arbitrary numbers that simply provide a relative IgA binding measure between the species and are used as the mean to generate a normal distribution of IgA binding for each species (with a standard deviation of one). For initial simulations, 30 samples were then generated with random pre-sort abundances for the ten species sampled from a log normal distribution.

To simulate an IgA-Seq experiment, each sample was assumed to have profiled 100,000 bacteria (a reasonable magnitude for 16S rRNA gene sequencing). For each sample, the 100,000 counts were assigned to one of the ten species based on the sample’s species abundances. Then, for each of the 100,000 bacteria in each sample, a value of IgA binding was generated at random from the normal distribution defined by its species mean. The distribution of IgA scores across all species was used to identify values that cut off the upper and lower tails of IgA binding and these were used to define the bacteria within the IgA+ and IgA- fractions respectively. The bacteria within the IgA+ and IgA- fractions for each sample were then used to generate the IgA+ and IgA- relative abundances and all other measures required for generating the various IgA-Seq scores.

A secondary simulation was performed to model the case where two groups are compared with a consistent difference in starting abundances between them. For this, the simulation was run using the same ten species as before but now considering 60 samples. Additionally, prior to generating bacterial IgA binding values, 0.5 was added to the abundance of species ten in half of the samples before renormalizing the abundances to sum to one. Samples with additional species ten were considered cases and the unaltered samples considered controls.

All stages of the IgA-Seq simulation were carried out using the function *simulateigaseq*, which is also part of the IgAScores package. All IgA-Seq indices were calculated as defined previously using the *igascores* function. For simulated data experiments, a value of 10^−6^ was used for pseudo counts when required as this was the same magnitude as the minimum non-zero abundance value observed in the simulated data. Correlations were assessed using the *cor* function in R [36]. Between group comparisons were performed using the *wilcox*.*test* function, adjusting p-values using *p*.*adjust*. Code for all simulation analyses can be found at https://github.com/microbialman/IgAScoresAnalyses.

### Bacterial culture

*Bacteroides thetaiotaomicron* VPI-5482 (*B. theta*) was cultured at 37°C in in brain heart infusion broth supplemented with 5 μg/ml hemin and 0.5 μg/ml vitamin K (BHIS) under anaerobic conditions (10% H2, 10% CO2, 80% N2). *Escherichia coli* LF82 was cultured aerobically at 37°C in Luria-Bertani (LB) medium. *Helicobacter hepaticus* ATCC 51449 was grown in tryptone soya broth (TSB) supplemented with 10% fetal calf serum and Skirrow Campylobacter supplements (Oxoid) under microaerophilic conditions (1-3% oxygen). To stain bacteria, pure overnight cultures were pelleted by centrifugation (8000 x *g*, 5 min, 4°C), washed in sterile phosphate buffered saline (PBS), and blocked with filter-sterilized 10% mouse serum (10 min, ice). The following monoclonal antibodies were used to stain bacteria in a 100 μl volume: mouse IgG2b anti-*E*.*coli* (1:100; Abcam clone 2D7/1) and mouse IgA anti-*B*.*theta* (1:100; clone 255.4[23]). To generate the 2-member community, the optical density (OD, 600 nm) and corresponding colony forming units (CFU) were determined for OD_600_ of 1: *E. coli* 8 × 10^7^ CFU; *B. theta* 1.2 × 10^8^ CFU. *Acutalibacter muris* KB18, *Akkermansia muciniphila* YL44, *Bacteroides caecimuris* I48, *Bifidobacterium animalis* subsp. *animalis* YL2, *Blautia coccoides* YL58, *Clostridium clostridioforme* YL32, *Clostridium innocuum* I46, *Enterococcus faecalis* KB1, *Flavonifractor plautii* YL31, *Lactobacillus* I49, *Muribaculum intestinale* YL27, and *Turicimonas muris* YL45 were used to generate stably colonized MM12 mice[37].

### Mouse strains

All experiments were conducted in accordance with the UK Scientific Procedures Act (1986) under a Project License (PPL) authorized by the UK Home Office. Animals were housed in accredited animal facilities at the University of Oxford and provided with sterile water, food ad libitum, and environmental enrichment. Both males and females were used in experiments after 8 weeks of age. Specific pathogen-free (SPF) C57BL/6J and C57BL/6J Rag1^−/−^mice were routinely tested and negative for *Helicobacter* spp. and other known intestinal pathogens. Germ-free (GF) C57BL/6J mice were maintained in sterile flexible film isolators and routinely screened for bacterial contamination by culturing, Gram stain, and 16S PCR. GF C57BL/6J mice were stably colonized with a defined consortium of 12 bacterial members, see below, of murine intestinal flora to generate a sDMDMm2[26, 37] (MM12) colony that was maintained in sterile flexible film isolators. All GF and MM12 mice were maintained on sterile food and water.

### *Helicobacter hepaticus* infection

To induce colitis using *Helicobacter hepaticus* (*Hh*), NCI-Frederick isolate 1A strain 51449 was grown as previously described [38]. Littermate control mice with MM12 flora were randomly assigned to groups. Mice were fed 1.0×10^8^ CFU by oral gavage on day 0 and 1. On day 0 and 7, 1mg IL-10 receptor blocking antibody (clone 1B1.2) was delivered by intraperitoneal injection. Control mice were housed in a separate flexible film sterile isolator and were given IL-10 receptor blocking antibody in the absence of *Hh* infection.

Animals were routinely monitored for weight loss and onset of diarrhea. Mice were sacrificed at day 14 and processed aseptically in a class II microbiological safety cabinet. Colon contents were collected for IgA-Seq and lipocalin-2 measurements (EMLCN2, Life Technologies) and stored at – 80°C. Formalin-fixed paraffin-embedded cross-sections of proximal, middle, and distal colon were stained with hematoxylin and eosin to assess histopathology. Sections were examined for hallmarks of inflammation: epithelial hyperplasia, goblet cell depletion, leukocyte infiltration, crypt abscess formation, submucosal leukocyte infiltration, and interstitial edema.

### Lipocalin-2 enzyme-linked immunosorbent assay

To prepare intestinal supernatants, 50-100 mg of colon contents were added to screw-capped O-ring tubes filled with 1 ml of cOmplete ULTRA EDTA-free protease inhibitor (Roche). Samples were homogenized on the TissueLyser II (Qiagen) for 10 minutes at 30 Hz and debris separated by centrifugation (10,000 x *g* for 5 min at 4°C). Supernatant was carefully collected and either used immediately or stored at −20°C. Lipocalin-2 was quantified using a mouse LCN2 enzyme-linked immunosorbent assay (ELISA) kit (ThermoScientific) according to manufacturer’s instructions. Each sample was run in duplicate and measurements reported as ng of lipocalin per gram of sample

### Preparation of samples for bacterial flow cytometry

Only sterile reagents were used to process samples. Frozen stool and intestinal contents (10-50 mg of sample) were rehydrated in 1 ml of PBS and suspended by vortexing. Large debris were separated by centrifugation (50 x *g*, 15 min, 4°C) and the clarified supernatant was passed through a 70 μm filter. A 500 μl aliquot of the suspension was transferred into a new 1.5ml microcentrifuge tube and an additional 1 ml of PBS added. Bacteria were pelleted (8000 x *g*, 5 min, 4°C) and blocked with 100 μl of 10% rat serum (10 min, ice). An additional 1 ml of PBS was added to the blocking suspension and centrifuged, and pelleted bacteria were washed with 1 ml of FACS buffer (PBS + 1% bovine serum albumin). Following centrifugation, the resulting pellet was stained with PE-conjugated rat anti-mouse IgA (1:100, eBioscience clone mA-6E1) in a 100 μl volume (30 min, covered on ice). Clone mA-6E1 was titrated testing concentrations of 1:10, 1:20, 1:50, 1:100, 1:200, 1:500, and 1:1000. Samples were washed twice with FACS buffer and resuspended in 1 ml buffer containing SYBR Green I nucleic acid gel stain (1:400,000, Life Technologies) in the dark at room temperature. The optimal concentration of SYBR was determined by titration testing a range from 1:100,000 to 1:500,000. Samples were then placed in a covered ice box and immediately acquired on the flow cytometer (LSRFortessa X20 and FACSAria III, BD Biosciences).

To validate the IgA-Seq protocol, 1 ml of an overnight *B. theta* culture was first stained with mouse IgA clone 255.4 (25 min, ice), washed with 1 ml PBS, followed by staining with PE-conjugated rat anti-mouse IgA (1:100, eBioscience clone mA-6E1). Stained bacteria were washed three times in FACS buffer before resuspension in 1 ml FACS buffer. A 500 μl aliquot of IgA-coated *B. theta* was spiked into a faecal suspension prepared from specific pathogen-free (SPF) C57BL/6J mice. SYBR was added to the final mixture.

### Fluorescence Activated Cell Sorting (FACS) of bacterial populations

Samples were sorted using a FACSAria III (BD Biosciences) instrument with external aerosol management. Both sample and collection tubes were maintained at 4°C. Sterile, preservative-free PBS without calcium or magnesium (Invitrogen) was used for sheath fluid and was autoclaved prior to sorting. The flow cytometer was sterilized according to the manufacturer’s Aseptic Sort protocol to decontaminate the entire sheath and sample paths. In addition, both before and after sorting, freshly made 10% bleach was run for 20 min on max flow rate followed by sterile PBS for 10 min. Sheath fluid from the droplet stream was collected prior to sorting as a control. Sample Line Backflush was run between samples to prevent carryover.

For finer resolution of small particles including bacteria, the neutral density filter was removed and threshold set to 200 for side scatter (SSC). Drop delay values were determined with the Accudrop feature and were updated with each new sort setup. To sort and culture live *B. theta*, various nozzle sizes (70, 85, 100 micron) and sheath pressures (20, 30, 45, 70 psi) were tested. A 10 × 10 grid was programmed into the automated cell deposition unit (ACDU) stage, and *B. theta* suspended in PBS was sorted in Single Cell precision mode onto BHIS agar plates for enumeration of colony forming units (CFU). Nozzle sizes and sheath pressures were also tested for sort purity of IgA+ and IgA- populations of faecal bacteria.

For stool and intestinal samples, the SYBR nucleic acid stain (FITC channel) helped to differentiate bacteria from autofluorescent debris (BV421 or PerCP Cy5.5 channels). Samples were diluted in FACS buffer containing SYBR in order to achieve an event rate of 5000-6000 events per second. To begin, 100,000 events were recorded from the initial sample (pre-sort) in order to calculate the percentage of bacteria in the IgA+ and IgA- gates used in the probability-based scores. Gates were then drawn to delineate bacterial populations that were bound with IgA (IgA+) or unbound (IgA-). The 4-way Purity precision mode was used to capture 400,000 events for each IgA+ or IgA- fraction. Fractions were collected into autoclaved 2 ml screw cap tubes (ThermoFisher Scientific) pre-coated with 200 μl FACS buffer. Sort purity of each fraction was recorded from 100 events, and DNA was immediately extracted from the remainder. BD FACSDiva and Flowjo software (TreeStar) were used for instrument setup and data analysis.

### Magnetic Activated Cell Separation (MACS)

As a comparison with FACS, IgA-stained samples were incubated for 30 min at 4°C with anti-PE microbeads (1:50, Miltenyi) in MACS buffer (0.5% bovine serum albumin and 2 mM EDTA in PBS). Microbeads were titrated by testing concentrations of 1:5, 1:20, 1:50, 1:100, 1:200, 1:500. Cells were pelleted by centrifugation (10,000 x g, 5 min, 4°C) and washed twice in MACS buffer before resuspension in 1 ml. LS columns (Miltenyi) were placed on a magnetic stand (Miltenyi) and pre-washed with 10 ml MACS buffer before the sample was loaded. Unlabelled “MACS neg” cells were collected in 3 × 3 ml washes. The LS column was then removed from the magnet and the labelled “MACS pos” cells were collected in 6 ml MACS buffer using the LS column plunger. MACS fractions were then analysed for purity by flow cytometry.

### DNA extraction

Only sterile molecular-grade reagents and filter-sterilized buffers were used to extract DNA. Sorted fractions were centrifuged (16,000 x *g*, 20 min, 4°C) and the supernatant removed such that only 300 μl remained in the tube. For unsorted (pre-sort) samples, 100 ul was transferred into an autoclaved 2 ml screw cap tube containing 200 μl FACS buffer. The following was then added to each tube: 250 μl autoclaved 0.1 mm zirconia/silica beads (Biospec), 300 μl Lysis buffer (200 mM NaCl; 200 mM Tris; 20 mM EDTA; pH 8), 200 μl 20% SDS, and 500 μl of phenol:chloroform:isoamylalcohol (PCI 25:24:1, pH 7.9, Sigma). Samples were bead beaten (6,500 rpm for 4 min, 3 min on ice, repeat 2x, Precellys tissue homogeniser), then centrifuged (6,000 x *g*, 10 min, 4°C) to separate phases. The aqueous phase was transferred into 5PRIME Phase Lock Gel Light tubes (Quantabio) and an equal volume of PCI added. After mixing by inversion, samples were centrifuged (16,000 x *g*, 10 min) and the aqueous phase transferred to a new Eppendorf tube. DNA was precipitated with 1/10 volume 3M NaOAc (pH 5.5) and 1 volume of isopropanol overnight at −20°C. After precipitation, DNA was pelleted (16,000 x *g*, 30 min, 4°C) and the supernatant carefully removed. The pellet was washed in 500 μl 70% ethanol, centrifuged (16,000 x *g*, 10 min, 4°C), and the wash repeated again. Residual ethanol was removed from pelleted DNA and the tube placed on a 37°C heat block until the pellet was completely dry. Cleaned DNA pellets were resuspended in 30 μl TE buffer at 50°C for 30 min, then stored at −20°C until 16S rRNA gene library preparation.

### 16S rRNA gene sequencing

The V4 region of 16S ribosomal RNA gene was amplified using barcoded primer pairs[39, 40]. Amplicons were generated in 50 μl PCR reactions: template DNA (2 μl), 10 μM 515F primer (1 μl), 10 μM 806R (1 μl), 5PRIME HotMasterMix (20 μl), and water (26 μl). Water was added instead of DNA for no-template controls. Thermocycler conditions: 94°C for 3 min; 35 cycles of 94°C for 45 s, 50°C for 60 s, and 72°C for 90 s; 72°C for 10 min. Amplicons were cleaned (UltraClean PCR Clean-Up Kit, Qiagen) and quantified (Quant-iT dsDNA high-sensitivity kit, ThermoFisher) in triplicate before pooling. The pooled library was precipitated with 2 volumes of 100% ethanol and a final concentration of 0.2M NaCl (45 min, ice). The supernatant was removed following centrifugation (7800 x *g*, 40 min, 4°C) and the pellet washed with 1 volume of 70% ethanol. The cleaned library was pelleted (7800 x *g*, 20 min, 4°C) and air dried to remove residual ethanol before resuspending in 100 μl DNAase-free water. Gel clean-up was performed (QIAquick Gel Extraction Kit, Qiagen) and the final library was sequenced on the MiSeq (Illumina) platform to generate 2 × 250 base pair paired-end reads.

All 16S rRNA gene sequencing data was quality checked using FastQC and processed using DADA2 within the NGSKit pipeline (https://github.com/nickilott/NGSKit) to generate an amplicon sequence variant (ASV) table[41]. DADA2 was used to assigned taxonomy to ASVs using a database built from a combination of both the RefSeq and RDP databases[42]. ASV counts were then summarised at the species level and all analyses carried out on these relative abundances (summing to one within a sample).

### IgA-Seq in a mouse model of colitis

ASV counts were generated from 16S rRNA gene sequences as described above, producing 54091±7970 (mean±SD) ASV counts per sample (excluding the blank control which had 5119 counts). These were summarised at the species level. Given the samples contained a known MM12 microbiota, the presence of unexpected species in samples and species observed in a sequencing control blank were used to find an abundance threshold that removed the majority of contaminants. Non-metric multi-dimensional scaling was used to visualise the relationship between samples from each mouse in each fraction using the *metaMDS* function from the vegan R package[43]. Comparisons of IgA-Seq data were performed between the four anti-IL10R only controls and the four *Hh* treated mice for the Palm, Kau and the Probability Ratio scores. These were generated using the *igascores* function as for the simulated data. A pseudo count of 10^−3^ was added where necessary as this was the magnitude of the lowest non-zero abundance value observed in the data.

Scores were compared between the colitic and control groups for taxa where there were at least three samples scored in both groups (taxa were scored NA if they had zero abundance in both the positive and negative fractions). Given the small sample size of each comparison significant differences were determined by carrying out all possible permutations of group labels and observing the difference in group means for each taxon for each score. The true difference in means was compared to the differences across permutations to derive an exact p-value, considering significance where p<0.1. This more permissive threshold was used given the small sample size (min three per group). To generate comparable effect sizes between scores, which are on different scales, the difference in means between groups was normalised to the standard deviations within the groups using the strictly standardised mean difference, calculated using the *ssmd* function from the *phenoDist* R package [44]. The species level data for these analyses are made available within the *IgAScores* package and all code for the analyses are available at https://github.com/microbialman/IgAScoresAnalyses.

### Analysis of Huus et al data

Sample metadata, IgA+ and IgA- fraction sizes, and the pre-processed taxon count table used in the original study were provided by the authors. The counts table was filtered to remove low abundance taxa using code published with the original study and then subset to just the samples for which the fraction sizes were available. The Kau Index and Probability Ratio scores were then generated using the *igascores* function as previously. For each taxon, a two-way ANOVA was performed, using the *aov* function in R, to determine if there was a difference in IgA binding score between diet treatments or the interaction of diet and mouse age. This was carried out for both the Kau Index and Probability Ratio, considering a taxon significant if either diet or the diet-age interaction had a nominal p<0.05. Taxa were only tested that had scores (i.e. were not absent in both the IgA+ and IgA- fractions) in at least six samples spread across all age points and both conditions. The magnitude of effects was estimated as the difference in mean scores between MAL and CON for each taxon and was calculated at each of the three ages individually. Mean differences were normalised by the standard deviation within groups, using the strictly standardised mean difference as for the colitis analysis (to make the values comparable between the Kau index and Probability Ratio). Coefficient of variation was calculated within each taxon within each experimental group (age and diet combined) for each scoring method and compared using a Mann-Whitney test. All code for the replication analysis can be found at https://github.com/microbialman/IgAScoresAnalyses.

## Declarations

### Ethics approval and consent to participate

All animal experiments were conducted in accordance with the UK Scientific Procedures Act (1986) under a Project License (PPL) authorized by the UK Home Office.

### Consent for publication

Not applicable

### Availability of data and materials

All code for the analyses in this study are available at https://github.com/microbialman/IgAScoresAnalyses. The IgAScores package is available at *https://github.com/microbialman/IgAScores* and also includes a pre-processed version of the data used in the analyses of the colitis model. The raw 16S rRNA gene sequencing data for the colitis experiment are available at the European Nucleotide Archive under study accession PRJEB39814 (https://www.ebi.ac.uk/ena/browser/view/PRJEB39814).

### Authors’ contributions

MAJ, CP, FP, LHL contributed to study design. MAJ,NEI,LHL contributed to data analyses. CP,ANH,JW,AJM,LHL contributed to experimental work. MAJ,KEH,BF contributed to replication analyses. MAJ,LHL authored the manuscript with inputs from all authors.

## Acknowledgements

The authors are grateful to Philip Ahern (Cleveland Clinic) and Jeffrey Gordon (Washington University) for providing the anti-*B. theta* antibody. We thank the Oxford Genomics Centre at the Wellcome Trust Centre for Human Genetics and the Genome Analysis Core at the Mayo Clinic for sequencing data generation. We also thank Elizete Araujo and members of the Oxford Centre for Microbiome Studies (OCMS) at the Kennedy Institute of Rheumatology for experimental support.

## References

1. Sommer F, Bäckhed F. The gut microbiota — masters of host development and physiology. Nat Rev Microbiol. 2013;11:227–38. doi: 10.1038/nrmicro2974.

2. Macpherson AJ, McCoy KD, Johansen F-E, Brandtzaeg P. The immune geography of IgA induction and function. Mucosal Immunol. 2008;1:11–22. doi: 10.1038/mi.2007.6.

3. Gutzeit C, Magri G, Cerutti A. Intestinal IgA production and its role in host-microbe interaction. Immunol Rev. 2014;260:76–85. doi: 10.1111/imr.12189.

4. Phalipon A, Cardona A, Kraehenbuhl JP, Edelman L, Sansonetti PJ, Corthésy B. Secretory component: a new role in secretory IgA-mediated immune exclusion in vivo. Immunity. 2002;17:107–15. doi: 10.1016/s1074-7613(02)00341-2.

5. Macpherson AJ, Köller Y, McCoy KD. The bilateral responsiveness between intestinal microbes and IgA. Trends Immunol. 2015;36:460–70. doi: 10.1016/j.it.2015.06.006.

6. Donaldson GP, Ladinsky MS, Yu KB, Sanders JG, Yoo BB, Chou W-C, et al. Gut microbiota utilize immunoglobulin A for mucosal colonization. Science (80-). 2018;360:795–800. doi: 10.1126/science.aaq0926.

7. Ansaldo E, Slayden LC, Ching KL, Koch MA, Wolf NK, Plichta DR, et al. Akkermansia muciniphila induces intestinal adaptive immune responses during homeostasis. Science (80-). 2019;364:1179–84. doi: 10.1126/science.aaw7479.

8. Moor K, Diard M, Sellin ME, Felmy B, Wotzka SY, Toska A, et al. High-avidity IgA protects the intestine by enchaining growing bacteria. Nature. 2017;544:498–502. doi: 10.1038/nature22058.

9. Mantis NJ, Rol N, Corthésy B. Secretory IgA’s complex roles in immunity and mucosal homeostasis in the gut. Mucosal Immunol. 2011;4:603–11. doi: 10.1038/mi.2011.41.

10. Grasset EK, Chorny A, Casas-Recasens S, Gutzeit C, Bongers G, Thomsen I, et al. Gut T cell–independent IgA responses to commensal bacteria require engagement of the TACI receptor on B cells. Sci Immunol. 2020;5:eaat7117. doi: 10.1126/sciimmunol.aat7117.

11. Macpherson AJ, Gatto D, Sainsbury E, Harriman GR, Hengartner H, Zinkernagel RM. A primitive T cell-independent mechanism of intestinal mucosal IgA responses to commensal bacteria. Science. 2000;288:2222–6. doi: 10.1126/science.288.5474.2222.

12. Bunker JJ, Flynn TM, Koval JC, Shaw DG, Meisel M, McDonald BD, et al. Innate and Adaptive Humoral Responses Coat Distinct Commensal Bacteria with Immunoglobulin A. Immunity. 2015;43:541–53. doi: 10.1016/j.immuni.2015.08.007.

13. Bunker JJ, Erickson SA, Flynn TM, Henry C, Koval JC, Meisel M, et al. Natural polyreactive IgA antibodies coat the intestinal microbiota. Science (80-). 2017;358.

14. Wei M, Shinkura R, doi Y, Maruya M, Fagarasan S, Honjo T. Mice carrying a knock-in mutation of Aicda resulting in a defect in somatic hypermutation have impaired gut homeostasis and compromised mucosal defense. Nat Immunol. 2011;12:264–70. doi: 10.1038/ni.1991.

15. Tsuruta T, Inoue R, Iwanaga T, Hara H, Yajima T. Development of a method for the identification of S-IgA-coated bacterial composition in mouse and human feces. Biosci Biotechnol Biochem. 2010;74:968–73. doi: 10.1271/bbb.90801.

16. Kullberg MC, Ward JM, Gorelick PL, Caspar P, Hieny S, Cheever A, et al. Helicobacter hepaticus Triggers Colitis in Specific-Pathogen-Free Interleukin-10 (IL-10)-Deficient Mice through an IL-12-and Gamma Interferon-Dependent Mechanism. Infect Immun. 1998;66:5157–66. doi: 10.1128/IAI.66.11.5157-5166.1998.

17. Palm NW, de Zoete MR, Cullen TW, Barry NA, Stefanowski J, Hao L, et al. Immunoglobulin A Coating Identifies Colitogenic Bacteria in Inflammatory Bowel Disease. Cell. 2014;158:1000–10. doi: 10.1016/j.cell.2014.08.006.

18. Kau AL, Planer JD, Liu J, Rao S, Yatsunenko T, Trehan I, et al. Functional characterization of IgA-targeted bacterial taxa from undernourished Malawian children that produce diet-dependent enteropathy. Sci Transl Med. 2015;7:276ra24–276ra24. doi: 10.1126/scitranslmed.aaa4877.

19. Viladomiu M, Kivolowitz C, Abdulhamid A, Dogan B, Victorio D, Castellanos JG, et al. IgA-coated E. Coli enriched in Crohn’s disease spondyloarthritis promote TH17-dependent inflammation. Sci Transl Med. 2017;9.

20. Huus KE, Bauer KC, Brown EM, Bozorgmehr T, Woodward SE, Serapio-Palacios A, et al. Commensal Bacteria Modulate Immunoglobulin A Binding in Response to Host Nutrition. Cell Host Microbe. 2020;:1–13. doi: 10.1016/j.chom.2020.03.012.

21. Kawamoto S, Maruya M, Kato LM, Suda W, Atarashi K, doi Y, et al. Foxp3+ T Cells Regulate Immunoglobulin A Selection and Facilitate Diversification of Bacterial Species Responsible for Immune Homeostasis. Immunity. 2014;41:152–65. doi: 10.1016/j.immuni.2014.05.016.

22. Catanzaro JR, Strauss JD, Bielecka A, Porto AF, Lobo FM, Urban A, et al. IgA-deficient humans exhibit gut microbiota dysbiosis despite secretion of compensatory IgM. Sci Rep. 2019;9:13574. doi: 10.1038/s41598-019-49923-2.

23. Peterson DA, McNulty NP, Guruge JL, Gordon JI. IgA Response to Symbiotic Bacteria as a Mediator of Gut Homeostasis. Cell Host Microbe. 2007;2:328–39. doi: 10.1016/j.chom.2007.09.013.

24. Kullberg MC, Jankovic D, Gorelick PL, Caspar P, Letterio JJ, Cheever AW, et al. Bacteria-triggered CD4+ T Regulatory Cells Suppress Helicobacter hepaticus–induced Colitis. J Exp Med. 2002;196:505–15. doi: 10.1084/jem.20020556.

25. Krausgruber T, Schiering C, Adelmann K, Harrison OJ, Chomka A, Pearson C, et al. T-bet is a key modulator of IL-23-driven pathogenic CD4+ T cell responses in the intestine. Nat Commun. 2016;7:11627. doi: 10.1038/ncomms11627.

26. Brugiroux S, Beutler M, Pfann C, Garzetti D, Ruscheweyh H-J, Ring D, et al. Genome-guided design of a defined mouse microbiota that confers colonization resistance against Salmonella enterica serovar Typhimurium. Nat Microbiol. 2017;2:16215. doi: 10.1038/nmicrobiol.2016.215.

27. Stallhofer J, Friedrich M, Konrad-Zerna A, Wetzke M, Lohse P, Glas J, et al. Lipocalin-2 Is a Disease Activity Marker in Inflammatory Bowel Disease Regulated by IL-17A, IL-22, and TNF-α and Modulated by IL23R Genotype Status. Inflamm Bowel Dis. 2015;21:1. doi: 10.1097/MIB.0000000000000515.

28. Palm NW, de Zoete MR, Cullen TW, Barry NA, Stefanowski J, Hao L, et al. Immunoglobulin A Coating Identifies Colitogenic Bacteria in Inflammatory Bowel Disease. Cell. 2014;158:1000–10. doi: 10.1016/j.cell.2014.08.006.

29. Lin R, Chen H, Shu W, Sun M, Fang L, Shi Y, et al. Clinical significance of soluble immunoglobulins A and G and their coated bacteria in feces of patients with inflammatory bowel disease. J Transl Med. 2018;16:359. doi: 10.1186/s12967-018-1723-0.

30. Castro-Dopico T, Dennison TW, Ferdinand JR, Mathews RJ, Fleming A, Clift D, et al. Anti-commensal IgG Drives Intestinal Inflammation and Type 17 Immunity in Ulcerative Colitis. Immunity. 2019;50:1099–1114.e10. doi: 10.1016/j.immuni.2019.02.006.

31. Simón-Soro Á, D’Auria G, Collado MC, Džunková M, Culshaw S, Mira A. Revealing microbial recognition by specific antibodies. BMC Microbiol. 2015;15:132. doi: 10.1186/s12866-015-0456-y.

32. Wilmore JR, Gaudette BT, Gomez Atria D, Hashemi T, Jones DD, Gardner CA, et al. Commensal Microbes Induce Serum IgA Responses that Protect against Polymicrobial Sepsis. Cell Host Microbe. 2018;23:302–311.e3. doi: 10.1016/j.chom.2018.01.005.

33. Acinas SG, Sarma-Rupavtarm R, Klepac-Ceraj V, Polz MF. PCR-Induced Sequence Artifacts and Bias: Insights from Comparison of Two 16S rRNA Clone Libraries Constructed from the Same Sample. Appl Environ Microbiol. 2005;71:8966–9. doi: 10.1128/AEM.71.12.8966-8969.2005.

34. Louca S, Doebeli M, Parfrey LW. Correcting for 16S rRNA gene copy numbers in microbiome surveys remains an unsolved problem. Microbiome. 2018;6:41. doi: 10.1186/s40168-018-0420-9.

35. Johnson JS, Spakowicz DJ, Hong B-Y, Petersen LM, Demkowicz P, Chen L, et al. Evaluation of 16S rRNA gene sequencing for species and strain-level microbiome analysis. Nat Commun. 2019;10:5029. doi: 10.1038/s41467-019-13036-1.

36. R Development Core Team. R: A language and environment for statistical computing. Vienna, Austria: R Foundation for Statistical Computing; 2009.

37. Li H, Limenitakis JP, Fuhrer T, Geuking MB, Lawson MA, Wyss M, et al. The outer mucus layer hosts a distinct intestinal microbial niche. Nat Commun. 2015;6:8292. doi: 10.1038/ncomms9292.

38. Maloy KJ, Salaun L, Cahill R, Dougan G, Saunders NJ, Powrie F. CD4+CD25+ TR Cells Suppress Innate Immune Pathology Through Cytokine-dependent Mechanisms. J Exp Med. 2003;197:111–9. doi: 10.1084/jem.20021345.

39. Caporaso JG, Lauber CL, Walters WA, Berg-Lyons D, Huntley J, Fierer N, et al. Ultra-high-throughput microbial community analysis on the Illumina HiSeq and MiSeq platforms. ISME J. 2012;6:1621–4. doi: 10.1038/ismej.2012.8.

40. Apprill A, McNally S, Parsons R, Weber L. Minor revision to V4 region SSU rRNA 806R gene primer greatly increases detection of SAR11 bacterioplankton. Aquat Microb Ecol. 2015;75:129–37. doi: 10.3354/ame01753.

41. Callahan BJ, McMurdie PJ, Rosen MJ, Han AW, Johnson AJA, Holmes SP. DADA2: High-resolution sample inference from Illumina amplicon data. Nat Methods. 2016;13:581–3. doi: 10.1038/nmeth.3869.

42. Alishum A. DADA2 formatted 16S rRNA gene sequences for both bacteria &amp; archaea. 2019. doi: 10.5281/ZENODO.2541239.

43. Dixon P. Vegan, a package of R functions for community ecology. J Veg Sci. 2003;14:927–30. doi: 10.1111/j.1654-1103.2003.tb02228.x.

44. Zhang X, Boutros M. A novel phenotypic dissimilarity method for image-based high-throughput screens. BMC Bioinformatics. 2013;14:336. doi: 10.1186/1471-2105-14-336.

