## Additional File 1: Supplements for "Accurate identification and quantification of commensal microbiota bound by host immunoglobulins"

**Figure S1.** Identification of IgA-coated bacterial populations by flow cytometry.
**Figure S2.** Sorting configuration enables sensitive sort control.
**Figure S3**. Between group comparisons of the Palm index in an IgA-Seq simulation.
**Figure S4.** Quality control to filter spurious taxa prior to analyses.
**Figure S5.** Comparison of IgA-Seq binding scores between groups in a mouse model of colitis.
**Figure S6.** Summary of the representation of the different scores and what questions they address.
**Table S1.** Mapping of bacterial strain names to their mappable identifiers in the 16S rRNA database used.

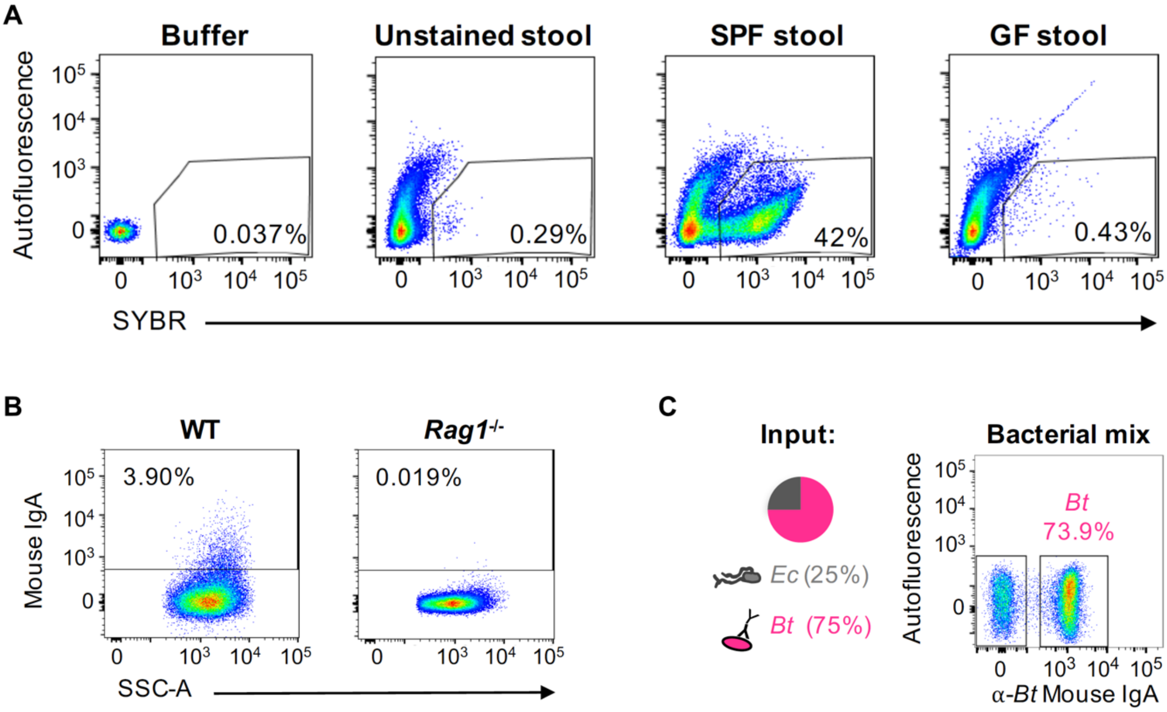


**Figure S1. Identification of IgA-coated bacterial populations by flow cytometry.** A) Gating SYBR DNA stain against autofluorescence (PerCP Cy5.5) differentiates faecal bacteria from debris. Filter-sterilised FACS buffer, unstained SPF C57BL/6J stool, and GF C57BL/6J stool as controls. Gated on events identified by forward (FSC) and side scatter (SSC) profiles. B) Percentage of endogenous faecal bacteria coated by IgA (PE, clone mA-6E1) in SPF C57BL/6J wild-type and *Rag1*$^{-/-}$ mice. Gated on SYBR+ cells. C) Artificial 2-member community of mouse IgG-coated E. coli (*Ec*) and mouse IgA-coated *B. theta* (*Bt*). Anti-mouse IgA (PE, clone mA-6E1) binds IgA+ *Bt* (α-*Bt* mouse IgA). Gated on SYBR+ cells.

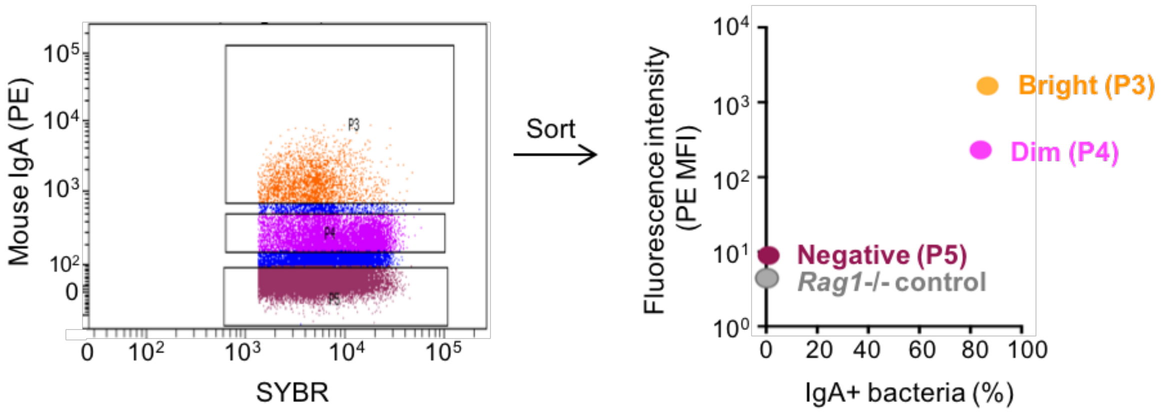


**Figure S2. Sorting configuration enables sensitive sort control.** Faecal bacteria from SPF C57BL/6 mice stained with anti-mouse IgA as in Figure 2. Sorting IgA$^{bright}$, IgA$^{dim}$ , and IgA- fractions with benchmarked protocol. Percentage of IgA+ bacteria and mean fluorescence intensity (MFI) of PE anti-mouse IgA for each collected fraction, including a *Rag1*$^{-/-}$ control.

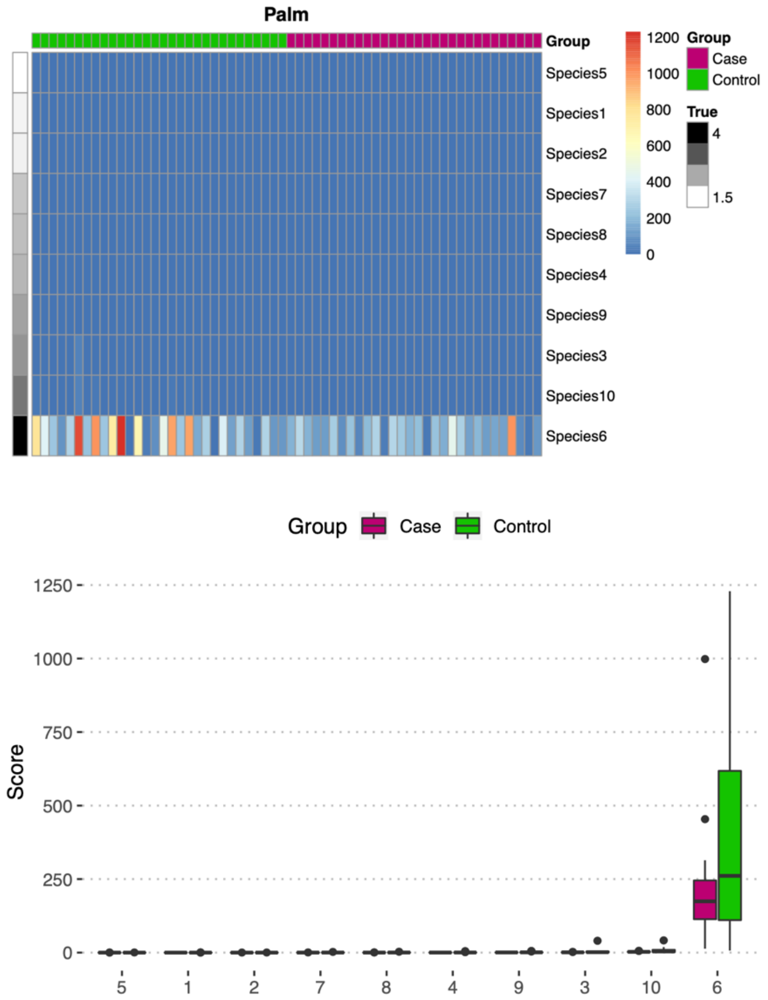


**Figure S3. Between group comparisons of the Palm index in an IgA-Seq simulation.** Plots are equivalent to those in Figure 5 for the Kau index and Probability Ratio. No significant differences were observed between groups in the Palm scores but it was only able to resolve the highly bound Species 6, where there was a much higher variability in the control group.

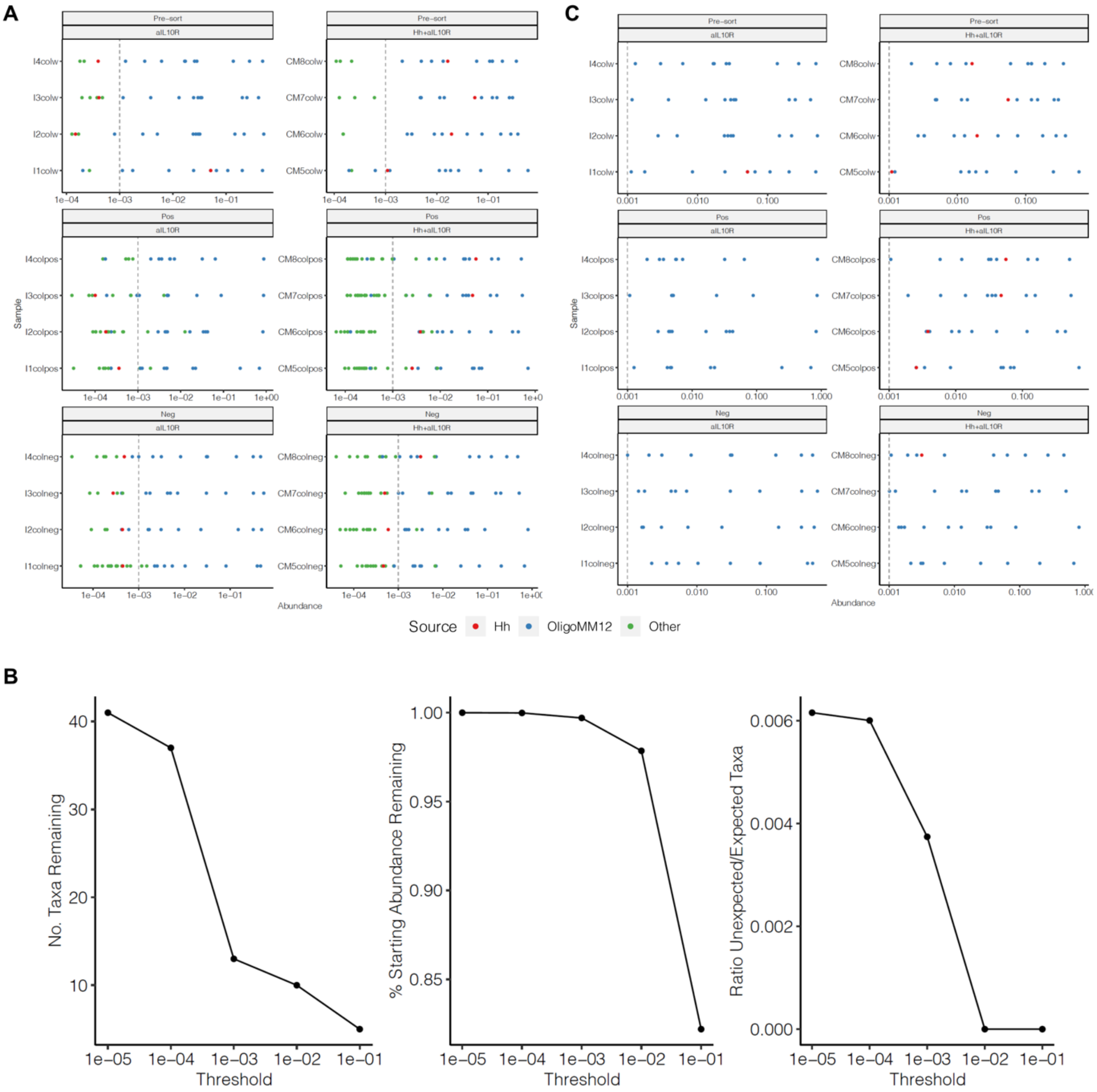


**Figure S4. Quality control to filter spurious taxa prior to analyses.** A) ASVs detected in samples after removing unexpected ASVs observed in the blank control. Remaining unexpected taxa are low abundance as shown in green. B) Plots used to select an appropriate cut-off for filtering out low-abundance taxa, the different thresholds considered are shown on the x-axis. The aim was to find a threshold that reduced the number of ASVs but retained the majority of abundance. C) Plot as in A after screening out low abundant ASVs and ASVs not observed in the pre-sort sample (in the IgA+ and IgA- fractions).

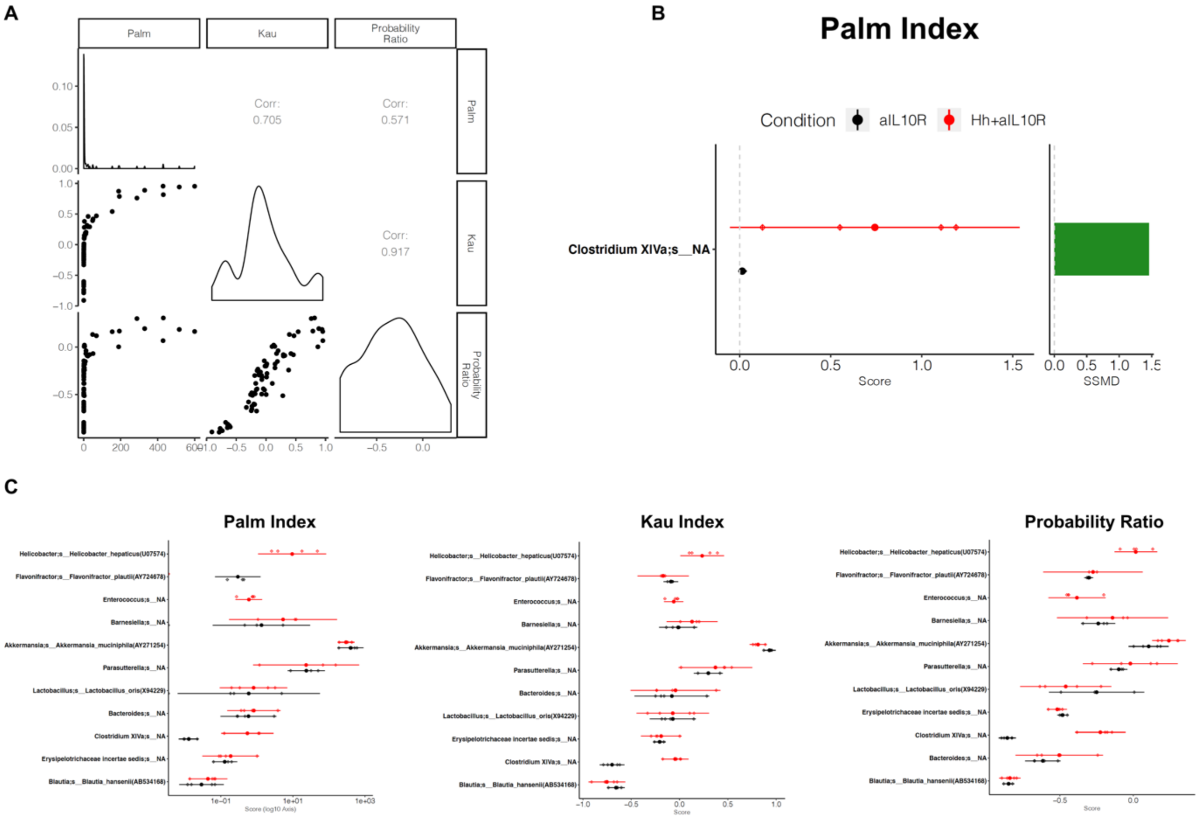


**Figure S5. Comparison of IgA-Seq binding scores between groups in a mouse model of colitis.** A) Pair-wise correlations between the scoring methods across all taxa from all mice, plots on the diagonal show the distribution of the scores. All correlations were significant (Pearson correlation p<0.001). B) Taxa with significantly different IgA-Scores in the aIL10R and Hh+aIL10R groups when using the Palm index, as for the other indices in Figure 6G. C) Plots showing the scores for all of the observed taxa in the MM12 comparisons using each of the Palm, Kau and Probability Ratio scores. Scores only observed in one group only have one colour (groups coloured as in B).

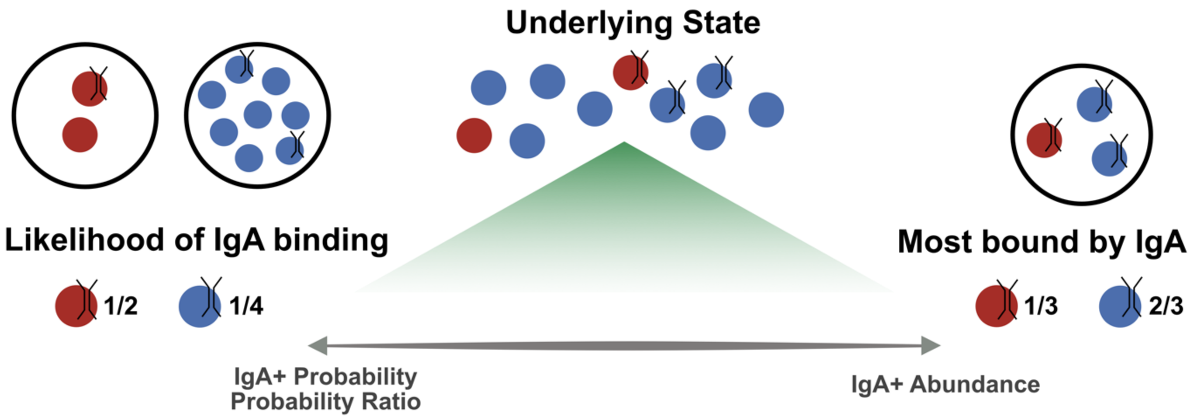


**Figure S6. Summary of the representation of the different scores and what questions they address.**

**Table S1. Mapping of bacterial strain names to their mappable identifiers in the 16S rRNA database used.**

| Strain Name | Species identifier in the DADA2 processed data |
|---|---|
| *Acutalibacter muris* KB18 | Clostridium XlVa;s__NA |
| *Akkermansia muciniphila* YL44 | Akkermansia;s__Akkermansia_muciniphila(AY271254) |
| *Bacteroides caecimuris* 148 | Bacteroides;s__NA |
| *Bifidobacterium animalis* subsp *animalis* YL2 | Not detectable in colon or stool |
| *Blautia coccoides* YL58 | Blautia;s__Blautia_hansenii(AB534168) |
| *Clostridium clostridioforme* YL32 | Clostridium XlVa;s__NA |
| *Clostridium innocuum* 146 | Erysipelotrichaceae incertae sedis;s__NA |
| *Enterococcus faecalis* KB1 | Enterococcus;s__NA |
| *Flavonifractor plautii* YL31 | Flavonifractor;s__Flavonifractor_plautii(AY724678) |
| *Lactobacillus reuteri* 149 | Lactobacillus;s__Lactobacillus_oris(X94229) |
| *Muribaculum intestinale* YL27 | Barnesiella;s__NA |
| *Turicimonas muris* YL45 | Parasutterella;s__NA |
| *Helicobacter hepaticus* | Helicobacter;s__Helicobacter_hepaticus(U07574) |
